# Anti-Immune Complex Antibodies are Elicited During Repeated Immunization with HIV Env Immunogens

**DOI:** 10.1101/2024.03.15.585257

**Authors:** Sharidan Brown, Aleksandar Antanasijevic, Leigh M. Sewall, Daniel Montiel Garcia, Philip J. M. Brouwer, Rogier W. Sanders, Andrew B. Ward

## Abstract

Vaccination strategies against HIV-1 aim to elicit broadly neutralizing antibodies (bnAbs) using prime-boost regimens with HIV envelope (Env) immunogens. Early antibody responses to easily accessible epitopes on these antigens are directed to non-neutralizing epitopes instead of bnAb epitopes. Autologous neutralizing antibody responses appear upon boosting once immunodominant epitopes are saturated. Here we report another type of antibody response that arises after repeated immunizations with HIV Env immunogens and present the structures of six anti-immune complexes discovered using polyclonal epitope mapping. The anti-immune complex antibodies target idiotopes composed of framework regions of antibodies bound to Env. This work sheds light on current vaccine development efforts for HIV, as well as for other pathogens, in which repeated exposure to antigen is required.

**One Sentence Summary:** Polyclonal epitope mapping reveals anti-immune complex antibodies which target idiotopes on antibodies bound to HIV Env.

## Introduction

Current vaccination strategies against HIV aim to elicit broadly neutralizing antibodies (bnAbs) against defined epitopes on the HIV envelope (Env) through efficient priming of specific B cells, the elimination of off target-epitope responses, and sustained boosting with antigens designed to guide particular patterns of somatic hypermutation associated with bnAbs (*1–3*). Many recent vaccine development efforts aimed at inducing bnAbs, utilize stabilized and soluble forms of trimeric Env, including SOSIP immunogens (*4*). The antibody responses to these SOSIP immunogens have been extensively studied using conventional serological approaches and more recently, electron-microscopy based polyclonal epitope mapping (EMPEM). Together these studies have shown that antibodies (Abs) primarily target a non-neutralizing, immunodominant neoepitope at the Env base (*5*) followed by a number of strain specific responses to other epitopes, such as the C3/V5 epitope, as well as neoepitopes like the N611 glycan hole (*6–8*). We have also shown that a subset of Abs that recognize the base of gp41 can induce dissociation into protomers *in vivo* and these base antibody-protomer immune complexes trigger Ab responses to non-neutralizing internal epitopes on Env (*9*). Thus, EMPEM has provided a new window into, and structural understanding of, *in vivo* immune complex formation and its influence on subsequent B cell receptor (BCR) engagement and secondary Ab elicitation. In this study we describe yet another phenomenon, anti-idiotypic and anti-immune complex antibodies elicited after repeat vaccination with SOSIP immunogens.

Antibodies can possess their own set of epitopes within the variable region of the Fab, called idiotopes. Abs that target idiotopes are thus called anti-idiotypic antibodies. Immune network theory (*10*) proposed that these anti-idiotypic antibodies result in a web of interacting idiotypes that start with antibodies (Ab1) produced in response to the antigen. These antibodies induce the production of anti-idiotypic antibodies (Ab2) which stimulate anti-(anti-idiotypic) antibodies (Ab3), and so on. These anti-idiotypic antibodies can be classified into three different categories: 1) Ab2ɑ antibodies recognize idiotopes of Ab1 that are outside of the antigen binding site but are part of the variable region; 2) Ab2β antibodies recognize idiotopes within the antigen binding site of Ab1 and mimic the original antigen epitope; 3) Ab2γ antibodies bind close to the antigen binding site of Ab1 but do not completely overlap with it, yet still interfere with antigen binding.

Here, we report the structures of six Ab2ɑ anti-idiotypic antibodies discovered during polyclonal epitope mapping of Ab responses to recombinant HIV Env SOSIP immunogens. These anti-idiotypic Abs were elicited against Abs recognizing variable epitopes on SOSIP immunogens in both rabbits and rhesus macaques. Anti-idiotypic Abs appear after repeated antigen exposure during boosting immunizations and recognize idiotopes composed predominantly of framework regions.

## Results

Stabilized, trimeric HIV Env immunogens have emerged as a promising vaccine platform for the induction of neutralizing antibodies and many concepts are being tested in human clinical trials. Iterative cycles of trimer immunogen design, testing, and characterization of immune responses remains critical for further optimization of this platform. As such, EMPEM is playing an increasingly central role in this process. Here we use EMPEM to elucidate how repeated immunizations with Env trimers results in the generation of previously unappreciated anti-immune complex (anti-IC) antibodies.

### Anti-Immune Complex Antibodies are Elicited by SOSIP Immunogens in Rabbits

#### Class I Anti-Immune Complex Antibodies

Negative stain (ns)EMPEM was performed to assess the immunogenicity of a 16055.v8.3 SOSIP antigen in New Zealand White rabbits (Fig. S1A) using week 22 serum complexed with matched 16055 antigen (*11*). nsEMPEM analysis of rabbit r2463 revealed polyclonal antibodies (pAbs) targeting the base neoepitope and adjacent regions of gp41 within single immune complexes. As these Abs appeared to contact each other, it motivated a higher resolution investigation using cryoEMPEM.

Briefly, cryoEMPEM employs a data processing pipeline based on consecutive rounds of 3D classification with spherical masks positioned around the epitope-paratope interfaces targeted by pAbs to pull out structurally unique, high-resolution classes. For the samples in this study, spherical masks encompassing the epitope-paratope interface of the pAb bound to HIV Env and the anti-immune complex antibody were used to computationally isolate particles containing the anti-IC Ab. Iterative rounds of 3D classification with different masks yielded high-resolution maps of the immune complex containing antigen, the primary Ab, and the anti-IC Ab (Fig. 1A and Fig. S2).

**Fig. 1.**
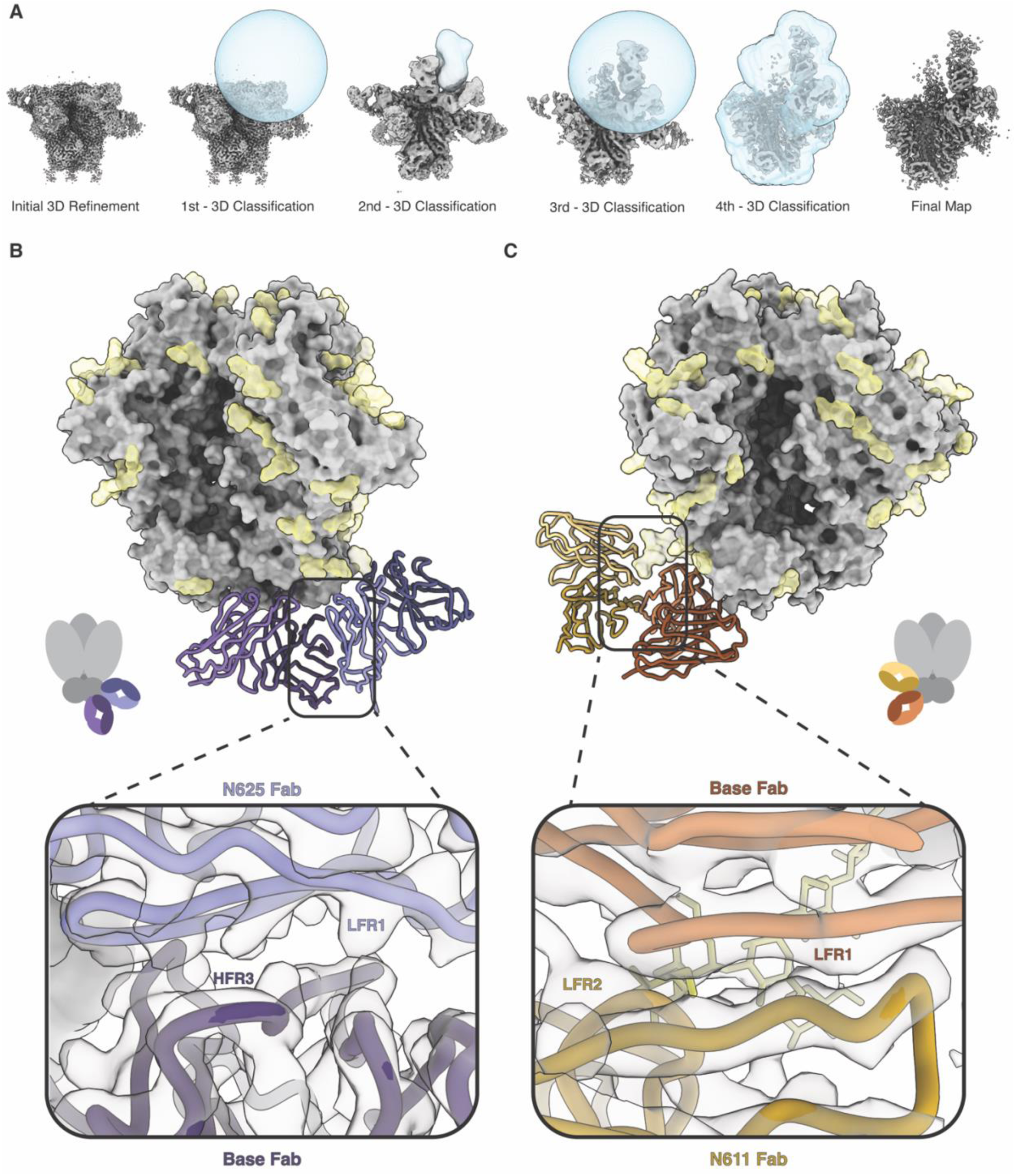
Class I Anti-Immune Complex Antibodies. CryoEMPEM was used to determine the structures of Class I anti-immune complex antibodies elicited during immunization with 16055 SOSIP. **A**) Consecutive rounds of 3D classification with spherical and antibody masks (blue) were used to isolate high-resolution classes of particles which contain the anti-immune complex antibody. The data processing pipeline with representative masks for each 3D classification step are shown, resulting in a final high-resolution map. **B)** A 3.3 Å resolution model of pAbs targeting the base neo-epitope and N625 glycan were resolved using cryoEMPEM. The SOSIP trimer is shown in gray with N-linked glycans depicted in yellow. The interface between these antibodies is shown. The light chain of the N625 Fab (periwinkle blue) contacts HFR3 of the base Fab (dark purple) through what appear to be aromatic residues in LFR1. **C)** A 3.8 Å resolution model of the same base Fab and a Fab targeting the N611 glycan are shown. There appears to be no specific contacts between the LFR2 of the N611 Fab (mustard yellow) and LFW1 of the base Fab (orange).

During cryoEMPEM data processing, we reconstructed a 3.3 Å resolution map with well-resolved epitope-paratope interfaces that included a pAb targeting the base epitope and a pAb bound to the N625 glycan (Fig. 1B). Analysis of the base and N625 pAbs revealed that these two Abs contacted one another along their framework regions (FR). However, the interactions appear to be limited, with the base pAb HFR3 contacting aromatic residues present in the LFR1 of the N625 pAb (Fig. 1B).

Additionally, from this dataset we isolated the same base pAb contacting a pAb targeting the N611 glycan at 3.8 Å resolution (Fig. 1C). In this interaction, the LFR1 of the base pAb makes minimal, non-specific contacts with LFR2 of the N611 pAb (Fig. 1C). The usage of this specific base pAb as a scaffold appears to be a preferred mode of binding for the N611 glycan targeting pAb since it is the predominant species observed during data processing. We also observed a small group of particles that contain the same N611 pAb as above but interact with a different base pAb. The resolution of this map was significantly lower, consistent with greater flexibility. The contacts between these Abs likely lead to greater stabilization of the complex.

Structural analysis of these two examples revealed that neither of the immune complexes isolated from 16055 r2463 qualify as anti-idiotypic antibodies because the antigen binding sites of these Abs contact the immunogen, not an idiotope on the adjacent Ab. However, these structures highlight the ability of the immune system to generate Abs with complementary surfaces that can tightly pack on the surface of an antigen. We thus define this type of antibody interaction as a Class I anti-immune complex antibody.

#### Class II Anti-Immune Complex Antibodies

A previous study evaluated whether trivalent or tetravalent SOSIP antigens were capable of eliciting bnAbs when delivered simultaneously or sequentially (*12*). In a monovalent control group, five rabbits were immunized with B41 SOSIP.v4.1(Fig. S1B). Epitope mapping was performed with week 22 serum complexed with B41 antigen. nsEMPEM analysis of rabbit r1646 revealed an anti-idiotypic antibody targeting an immune complex composed of B41 SOSIP and a pAb targeting the N241 glycan hole epitope.

CryoEMPEM processing of the B41 r1646 immune complex resulted in a 5.8 Å resolution map (Fig. 2A). Due to the low local resolution in the pAbs (5.0-5.5 Å), an initial model of a B41 SOSIP.v664 trimer and reference Fabs were docked into the reconstructed cryoEMPEM map without refinement. Resolution was sufficient to assign complementarity-determining regions (CDRs) and FRs. Analysis of the docked model revealed that the antigen binding pAb contacts the epitope comprising N241 glycan and surrounding C2 and C3 regions in B41 using HCDR1, HCDR3, and LCDR3 as well as contacts with the N88 glycan using HCDR2 and HFR3. The local resolution of the anti-idiotypic pAb is lower (5.5-6.0 Å), but we could discern the anti-idiotypic pAb contacts LFW1 of the N241 pAb using LCDR1. The pAb also contacts the N339 glycan on the B41 antigen using LCDR3, HCDR3 and HCDR2 (Fig. 2A). Unlike the Class I anti-IC Abs elicited against 16055 SOSIP, this anti-idiotypic antibody bound to a neoepitope comprised of both the N241 pAb and B41 SOSIP using its CDR loops. Therefore, we refer to this type of interaction as a Class II anti-immune complex antibody.

**Fig. 2.**
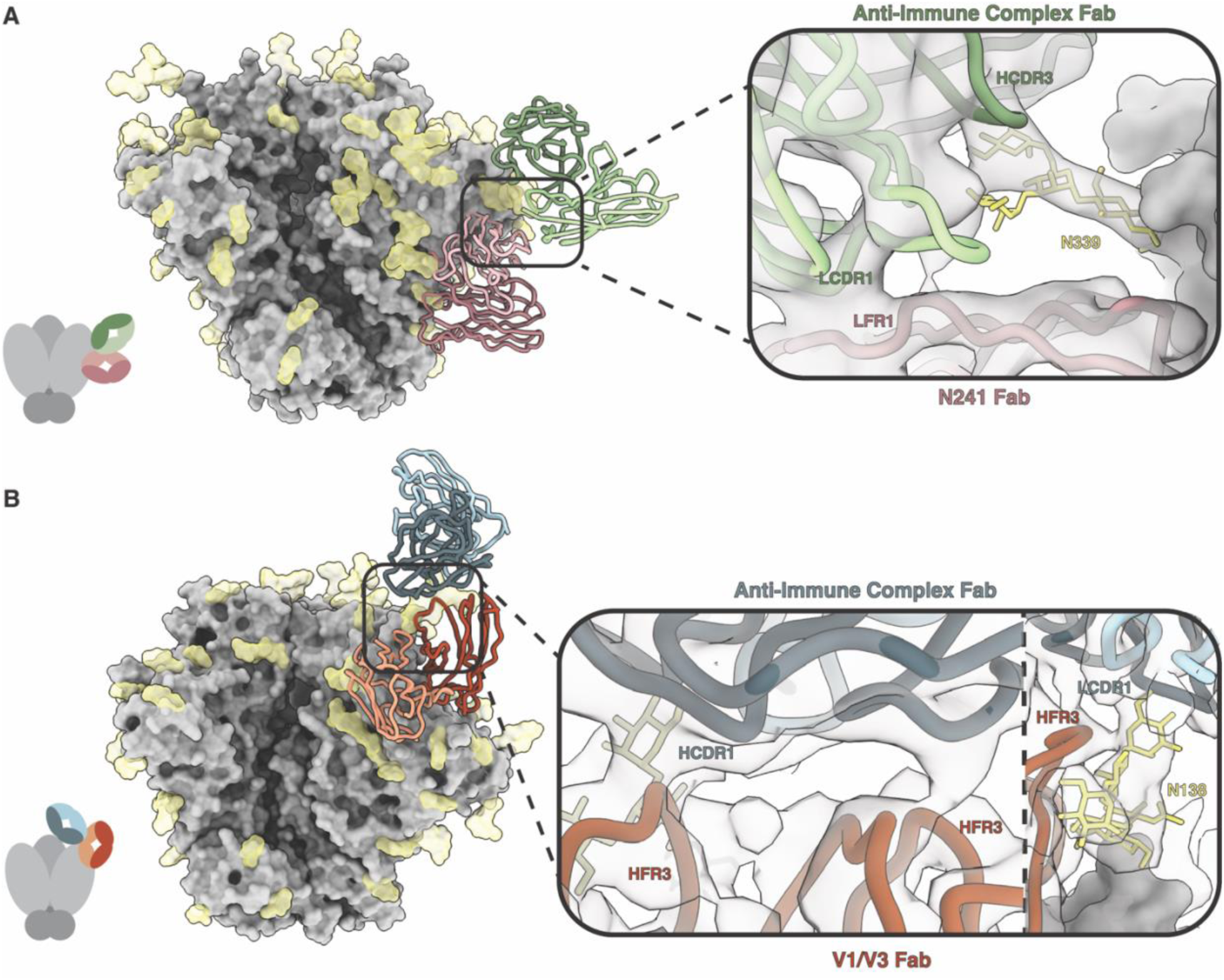
Class II Anti-Immune Complex Antibodies Contact Both Antigen and Antibody. CryoEMPEM was used to determine the structures of Class II anti-immune complex antibodies (anti-IC Abs) whose epitopes are composed of both antigen and antibody. **A)** An immunization with B41 SOSIP elicited an anti-IC Ab (green) that targets an antibody which recognizes the N241 glycan (pink). Due to low resolution (5.8 Å), rabbit anti-HIV Fabs (PDB ID: 6CJK) were docked into the map to visualize interactions between the two antibodies. The anti-IC Ab contacts the N339 glycan using HCDR2 and HCDR3 (green). It makes additional contacts with LFR1 of the N241 Fab (pale pink) using LCDR1 (light green). The N339 glycan is shown in yellow. **B)** An immunization with a chimeric CH505/BG505 SOSIP antigen elicited an anti-IC Ab (blue) that binds to a pAb targeting the V1/V3 loops (orange). This structure was resolved to 4.4 Å. The anti-IC Fab (dark blue) can be seen contacting HFR3 of the V1/V3 Fab (dark orange) using HCDR1. On the right side of the panel, the N138 glycan can be seen in yellow contacting LCDR1 of the anti-immune complex Fab (light blue) and HFR3 of the V1/V3 Fab (dark orange).

An additional immunization in rabbits was performed using a chimeric CH505/BG505 SOSIP.v8.1 antigen (Fig. S1C). Epitope mapping was performed with week 26 serum complexed with the chimeric antigen. nsEMPEM analysis of rabbit r2474 revealed a Class II anti-idiotypic antibody. CryoEMPEM analysis of the CH505/BG505 r2474 immune complex resulted in a 4.4 Å resolution map. The immune complex contained two Abs, one of which targeted the V1/V3 interface epitope on the antigen, and a second anti-idiotypic antibody that bound to an epitope comprised of V1/V3 pAb framework regions and the N138 glycan on the antigen (Fig. 2B). The anti-idiotypic antibody contacts the N138 glycan using both LCDR3 and HCDR3 and interacts with the V1/V3 pAb HFR2 and HFR3 using HCDR1 and HCDR2 (Fig. 2B).

Interestingly, we observed a third pAb bound to the trimer at the V2 apex in many of the same classes as the anti-IC Abs. A majority of particles possessed all three pAbs, resulting in densely packed Abs atop the SOSIP antigen (Fig. S5). The final reconstructed map showed some diffuse density for the V2 apex pAb, but it appears this additional Ab does not contact either the V1/V3 pAb or anti-IC Ab and therefore was not modeled. Our observations are consistent with pre-existing Ab responses to Env shaping ensuing responses.

#### Class III Anti-Immune Complex Antibodies

An additional rabbit from the 16055 SOSIP.v8.3 immunization study also appeared to elicit a different class of anti-idiotypic antibody when analyzed with nsEMPEM. CryoEMPEM of the 16055 r2464 immune complex yielded a 3.3 Å resolution map with well-resolved epitope-paratope interfaces for both pAbs. Analysis of the atomic model revealed two pAbs, one of which targets the V2 epitope on 16055 SOSIP and a second anti-idiotypic antibody that binds to an epitope entirely comprised of V2 pAb heavy chain framework regions (Fig. 3). For this reason, we define this canonical Ab2ɑ anti-idiotypic antibody interaction as a Class III anti-immune complex antibody. Upon further analysis, the anti-idiotypic antibody primarily uses LCDR3 and HCDR2 to make multiple shallow contacts within the FR1 and FR3 regions on the V2 pAb heavy chain (Fig. 3). Many of the contacts in the epitope-paratope interface appear to be between aromatic residues in the framework regions of the V2 Fab and CDR loops of the anti-IC Ab (Fig. S6). Extensive π-π stacking networks likely contribute to antibody binding and stabilization at this interface.

**Figure 3.**
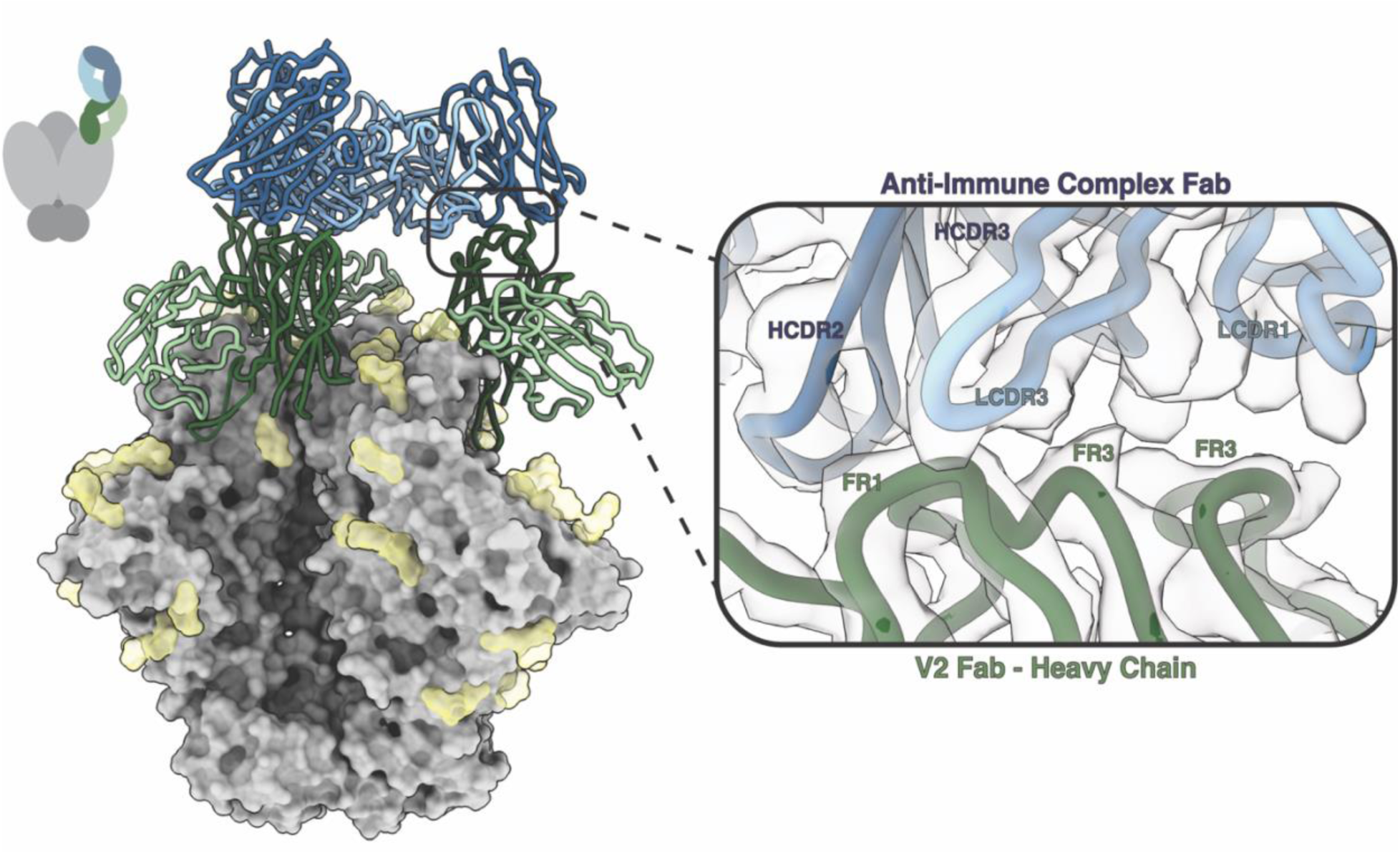
Class III Anti-Immune Complex Antibodies Have Epitopes Which Contain Antibody Framework Regions. An additional class of anti-immune complex antibody (anti-IC Ab) was elicited during vaccination with 16055 SOSIP. A 3.3 Å resolution map of a polyclonal antibody targeting the V2 epitope (green) and an anti-IC Ab (blue) was determined using cryoEMPEM. The anti-IC Fab contacts HFR1and HFR3 of the V2 Fab (dark green) using LCDR1 and LCDR3 (light blue) and HCDR3 (dark blue).

### BG505 SOSIP Immunization in Rhesus Macaques Can Elicit Anti-Immune Complex Antibodies

#### Class IV Anti-Immune Complex Antibodies

The elicitation of anti-IC Abs in rabbits has been observed for several different HIV Env SOSIP immunogens as shown above. In a previously published study, we analyzed non-human primate antibody responses (*13*). In this study, six rhesus macaques were immunized with BG505 SOSIP.v5.2 N241/N289 (Fig. S1D). Epitope mapping experiments were performed with week 38 serum complexed with the BG505 SOSIP antigen. Epitope analysis of Rh.33203 revealed an anti-idiotypic antibody targeting an immune complex composed of BG505 SOSIP and two pAbs, one of which targeted the gp120-interface epitope and the second the V5 loop.

Further analysis with cryoEMPEM produced a map of this immune complex at 4.6 Å resolution with moderate resolution in the SOSIP-V5 pAb and SOSIP-gp120-interface pAb epitope-paratope interfaces (Fig. 4). To improve the resolution of these two pAbs and better visualize contacts with antigen, individual models of the gp120-interface pAb (3.8 Å) and V5 pAb (4.0 Å) were isolated during data processing and combined in Chimera (*14*) to generate a starting model for the immune complex. The resolution at the anti-idiotypic antibody epitope-paratope interface was low (6-6.5 Å) and thus was refined only to a small degree.

**Figure 4.**
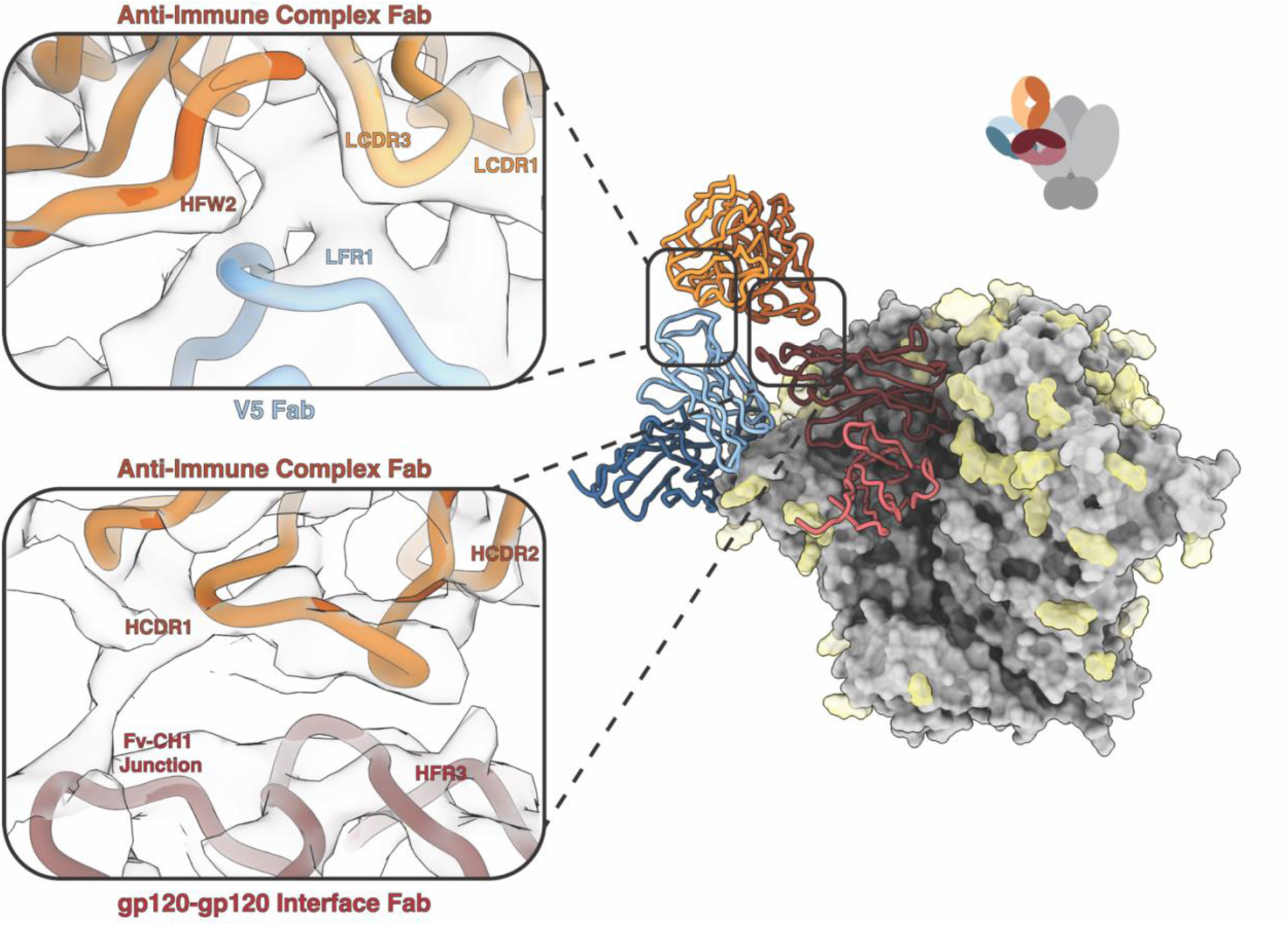
Class IV Anti-Immune Complex Antibodies Bind to Two Polyclonal Antibodies. CryoEMPEM was used to determine the structure of a Class IV anti-immune complex antibody (anti-IC Ab) whose epitope is composed of two separate pAbs. The 4.6 Å resolution map contains three pAbs, one of which targets the V5 epitope (blue), one which targets the gp120-interface epitope (red) and an anti-IC Ab (orange) which bridges the two. The local resolution at the epitope-paratope interface is low (6-6.5 Å) making it difficult to determine specific contacts. The top panel shows the interface between the V5 Fab and anti-IC Fab, with LCDR3 (light orange) and HFR2 and HCDR3 (orange) of the anti-IC Ab contacting LFR1 (light blue) of the V5 Fab. The bottom panel shows the interface between the gp120-interface Fab and anti-IC Fab. The anti-IC Fab contacts the Fv-CH_1_ junction and HFR3 of the gp120-gp120 interface Fab (brick red) using HCDR1 and HCDR2 (dark orange). There are no observable contacts between the anti-IC Ab and the SOSIP antigen.

The BG505 Rh33203 model contains three pAbs, one of which targets the gp120-interface epitope and a second antibody that binds to the V5 loop epitope. Interestingly, the third anti-idiotypic antibody binds to an epitope that bridges both the gp120-interface and V5 pAbs. It does not appear to interact with the SOSIP antigen, creating what we now refer to as a Class IV anti-immune complex antibody. Due to the low resolution at the anti-IC Ab epitope, it is difficult to determine specific contacts, but it appears that the anti-IC Ab contacts the V5 pAb LFR1 using LCDR3, HCDR3, and HFR2 (Fig. 4 left inset). At the gp120-interface pAb epitope, the anti-IC pAb contacts the Fv-CH_1_ junction using HCDR1 and contacts HFR3 using HCDR2 (Fig. 4 right inset). To our knowledge, this binding mechanism in which the anti-idiotypic antibody binds to an idiotope composed of two antibodies has not been previously described.

### Anti-Immune Complex Antibodies Are Elicited After Repeated Exposure to Antigen

To determine when anti-IC Abs are elicited during a vaccination regimen, nsEMPEM was performed using serum samples collected at a series of different time points. In the 16055 SOSIP (*11*) and chimeric CH505/BG505 SOSIP immunizations, no Ab responses were observed at the study start (W0) or two weeks after the first immunization (W2). Following the second immunization (W6) an Ab against the base neoepitope appeared in both studies (Fig. 5A-C). The B41 SOSIP immunization (*12*) produced a base response after the first immunization (W4 and W6), but also developed an Ab against the N611 glycan hole at these time points (Fig. 5D). Importantly, the precursor Ab that is recognized by the anti-IC Ab was not observed in these early time points. Following the third immunization (W22) the anti-IC Ab appears alongside an expanded Ab repertoire against other epitopes. By extrapolation, it is anticipated that the precursor Ab preceded the anti-IC Ab between these timepoints for which we did not have serum to analyze. Longitudinal nsEMPEM also enabled us to assess how the pAb response in rhesus macaques developed in response to immunization with BG505 SOSIP (*13*). After the second immunization (W10) Ab responses to the base and N289 glycan epitopes were observed. Following the third immunization (W26) precursor V5 and gp120-interface Abs for anti-idiotypic antibody recognition appeared. After the fourth immunization (W38) the anti-idiotypic antibody appeared (Fig. 5E). Hence, our data demonstrate that anti-IC Abs are elicited after repeated exposure to antigen after the second or third boosting immunization, at which point primary Ab targets have been elicited and are circulating at detectable levels in serum.

**Figure 5.**
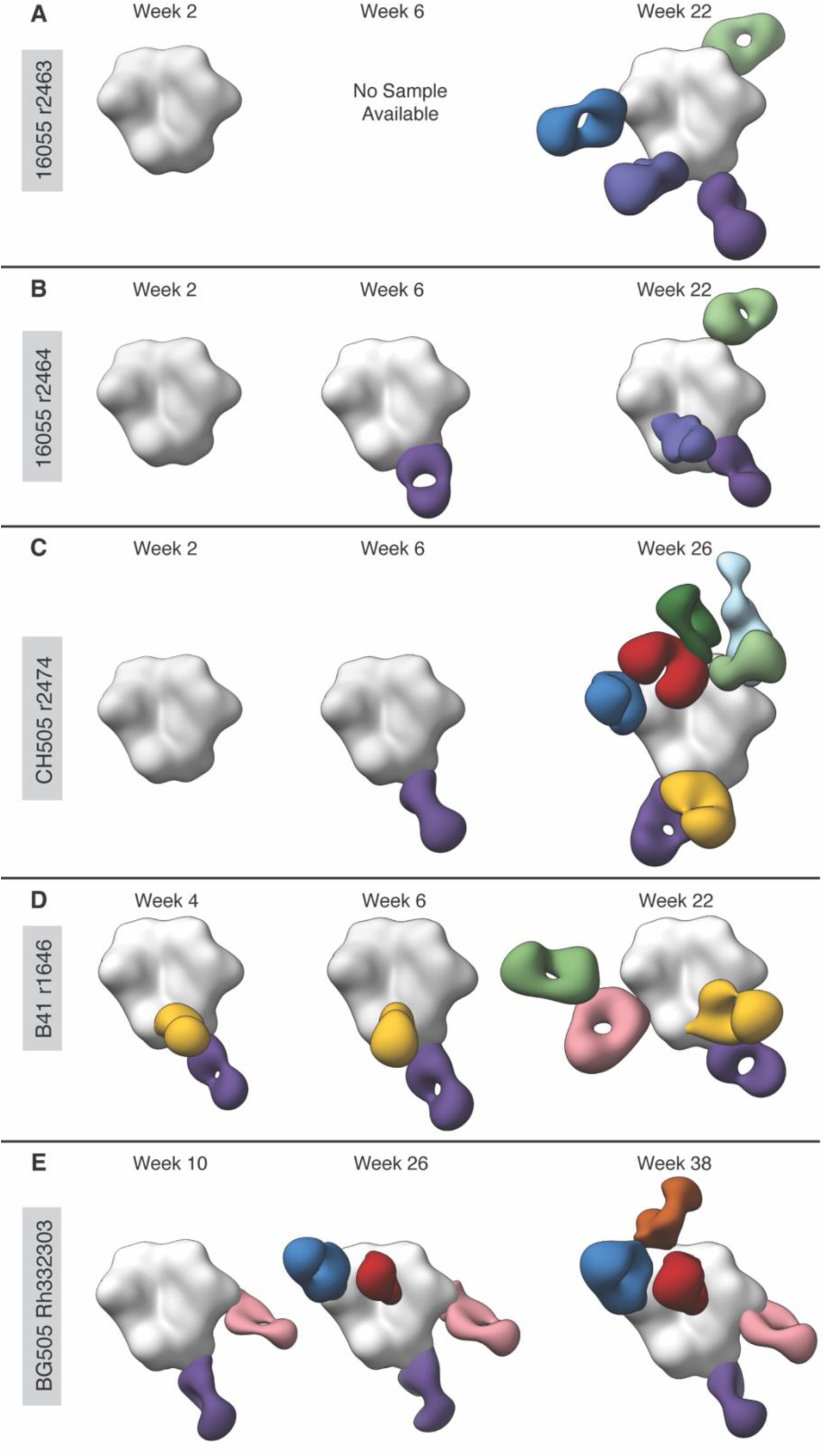
Longitudinal nsEMPEM Shows Anti-Immune Complex Antibodies Are Elicited After Repeated Exposure to Antigen. Longitudinal nsEMPEM was performed using serum samples from animals that elicited anti-immune complex antibodies (anti-IC Abs). No Ab responses were observed at the study start (W0). **A)** The 16055 r2463 samples show no Ab response after the first immunization (W2). By the third immunization (W22) the anti-IC Ab appears alongside an expanded Ab repertoire of base, N625 glycan, V5 loop, and V1/V2/V3 Abs. **B)** The 16055 r2464 samples also show no Ab response after the initial immunization (W2) and a single response to the base epitope after the second immunization (W6). After the third immunization (W22), Abs targeting the N625 glycan and V2 epitope appear. **C)** The CH505 r2474 samples show no Ab response after the initial immunization (W2) and a single response to the base epitope after the second immunization (W6). After the third immunization (W26) the anti-IC Ab (light blue) appears alongside an expanded Ab repertoire of base, N611 glycan, V5 loop, gp120-gp120 interface, V1/V3 and V2 apex Abs. **D)** The B41 r1646 samples show responses to the N611 glycan and base epitopes at after the first and second immunizations (W4 and W6). After the third immunization (W22) the N241 glycan epitope Ab and anti-IC Ab appear. **E)** The BG505 Rh.33203 samples show responses to the base and N289 epitopes after the second immunization (W10), and V5 loop and gp120-interface Abs after the third immunization (W26). After the fourth immunization (W38) the anti-IC Ab appears (orange).

### Anti-Immune Complex Antibody Paratopes Are Enriched with Aromatic Residues

The Class III 16055 2464 anti-IC Ab has a well resolved epitope-paratope interface, making it a strong candidate for automated atomic model building using ModelAngelo (*15*). This tool combines information from the cryoEM map, structural information, and predicted amino acid probabilities to objectively build atomic models into areas with unknown sequences. Using ModelAngelo, we generated predicted sequences for the anti-IC and V2 pAbs and used them to search a next generation sequencing (NGS) database of unpaired BCR sequences generated from peripheral blood mononuclear cells (PBMCs). We were able to identify sequences for the anti-IC heavy and light chains and V2 heavy chain that closely matched the EM map (Fig. S7). Unfortunately, the predicted V2 light chain did not pull any hits from our repertoire, thereby prohibiting experimental verification of the recovered Ab sequences.

Detailed inspection of reconstructed EM maps suggests that the HCDR3 and LCDR3 loops of the anti-IC Ab are enriched with aromatic residues as evidenced by density corresponding to aromatic side chains (Fig. S6 and S7). To test this hypothesis, we performed a frequency analysis to determine the likelihood of having aromatic residues at specific positions in these loops. Using an alignment of the NGS BCR sequences we calculated the amino acid probability distribution and identified sequence motifs based on amino acid properties (polarity, charge, hydrophobicity, aromaticity) for the anti-IC Ab (Fig. S8 and S9). LCDR3 contains two aromatic residues at positions 93 and 94 (IMGT numbering)(Fig. S8A). Based on our analysis, the frequency of an aromatic amino acid occurring at these positions is 26% and 70% respectively. Neither of the residues show significant amino acid frequency for Phe, indicating they are likely Tyr residues. Interestingly, Tyr94 makes a key contact with residue 82 on the V2 heavy chain, indicating this residue is likely conserved and essential for Ab binding. HCDR3 also contains two aromatic residues at positions 96 and 98. The frequency of an aromatic residue occurring at these positions is 25% and 55% respectively, with Tyr96 and Phe98 being the most likely identities (Fig. S8B). This Phe98 residue makes stabilizing contacts with the light chain, whereas Tyr96 makes a key interaction with an aromatic residue at position 64 in the V2 HC framework. As such, it appears both the light and heavy chains of the 16055 2464 anti-IC Ab are enriched for aromatic residues, one of which makes essential contacts with the V2 pAb, when compared to the BCR sequence repertoire from this rabbit.

### Anti-Immune Complex Antibodies Make Shallow Contacts with Shorter HCDR3 Loops

We observe that these anti-IC Abs contact their epitope using shorter, shallow HCDR3 loops compared to pAbs bound to Env. HCDR3 lengths were estimated using key anchor residues visible in the cryoEM maps which can only be assigned for maps below ∼4.0 Å local resolution, and thus were only able to be determined for 16055 r2464, 16055 r2463, and CH505 r2474 Abs. The average HCDR3 length for the Class II and Class III anti-IC Abs was 12.5 amino acids (n=2) and the average HCDR3 length for Abs bound to Env was 17.4 amino acids (n=5) (Fig. S10). Rabbits have an average HCDR3 length of 12 amino acids (*16*, *17*) which is consistent with our observed lengths for anti-IC Abs. Conversely, the pAbs bound to Env observed in these models are considerably longer than that average (17.4 aa).

## Discussion

Recently, studies investigating primary Ab responses on antigen presentation and secondary immune responses have shown that peripheral Abs can inhibit or enhance naïve B cell entry into germinal centers (*18–20*). Tas et al., showed that low affinity or low concentrations of circulating Abs enhance recruitment of B cells to germinal centers, whereas epitope-specific high affinity Abs elicited by HIV bnAb priming immunogens attenuate further epitope-specific B cell responses (*18*). Thus, pre-existing high affinity Abs can bias germinal center B cell activation and memory B cell selection by increasing or decreasing the activation threshold of B cells and directly masking epitopes on the antigen. After the initial priming immunization, low antigen availability in the germinal center promotes B cells that have high affinity for antigen and do not require high levels of somatic hypermutation. The formation of immune complexes with vaccine antigen can also result in multimerization, leading to increased avidity for the multimerized immune complex, resulting in the recruitment of B cells with low-affinity receptors into the germinal center (*19*). These B cells likely target immunodominant epitopes. Upon boosting, higher antigen availability and epitope masking of immunodominant epitopes promotes the evolution of subdominant responses in secondary germinal centers (*19*, *20*). This mechanism of inhibition or enhancement based on Ab affinity and availability can divert the immune response away from generating bnAbs to produce non-neutralizing Abs against immunodominant epitopes or neoepitopes on HIV. Here we show that Ab responses can be further diverted to epitopes presented on immune complexes of foreign antigen and host pAbs.

We speculate that after immunization with the initial priming immunogen, naïve B cells enter the germinal center and B cells with high affinity to Env proliferate and affinity mature to target immunodominant epitopes on HIV Env, such as the base neoepitope. After boosting with antigen, these pre-existing immunodominant Abs efficiently form immune complexes with antigen, resulting in epitope masking of subdominant epitopes. We believe that these immune complexes may be presented to follicular dendritic cells (FDCs) as multimers, increasing the apparent affinity of a B cell by avidity effects and enabling the recruitment of B cells with lower affinity into the germinal center. Due to the heavy glycosylation of HIV Env and epitope masking of the underlying peptide surface by the first series of immunodominant antibodies, the only epitopes available on these highly avid immune complexes that can be recognized by B cells are epitopes which are traditionally immunosilent (Fig. S11). The presence of pre-existing pAbs in these immune complexes may stabilize N-linked glycans or other flexible epitopes which are normally subdominant or immunologically silent. These pAbs have the ability to prime secondary Ab responses, leading to mature B cells that recognize epitopes which may include stabilized N-linked glycans to elicit the diverse range of anti-IC Abs we observe against framework idiotopes (Fig S11).

Our data demonstrate the BCRs must recognize immune complexes *in vivo* that are densely packed with pre-existing antibodies. Tight Ab packing has been observed for other antigens, namely to the internal repeat regions of the *Plasmodium falciparum* circumsporozoite protein (PfCSP). These anti-PfCSP Abs produce a spectrum of helical structures in which Fab is tightly packed against PfCSP. There are two important differences with these Abs compared to the Abs described here. First, because anti-PfCSP Abs bind repeat regions they are all identical, or homotypic, while Abs to adjacent epitopes on Env are heterotypic. Second, the anti-PfCSP Abs undergo somatic hypermutation outside the antigen paratope to form specific interactions with adjacent Abs, which increases binding avidity and results in increased protection against malaria infection (*21–23*). Recently, heterotypic antibody interactions have been described for *P. falciparum* cysteine-rich protective antigen (CyRPA). Three mAbs that target the same face of CyRPA were shown to synergistically interact with one another through lateral heterotypic interactions between framework regions, stabilizing the complex and slowing dissociation from CyRPA (*24*). In both of the examples above, it is evident that immune complexes are presented to BCR intact, clearly shaping subsequent BCR engagement.

Notably, the unique format of directly imaging unpurified immune complexes via EMPEM made our observations possible and illuminates how complex signals may be compressed in conventional serological assays such as ELISA. Further, because of the dependency on primary antigen binding Abs, conventional Ab generation via single cell or hybridoma methodologies are very unlikely to produce anti-IC antibodies. Antigen specific B cell sorting is also unlikely to generate anti-idiotypic antibodies as it would likely require two B cells to come into close proximity and overcome a huge steric hurdle. Conversely, our work demonstrates that cryoEMPEM can detect low abundance anti-IC Abs in polyclonal sera and characterize them in high resolution detail. The structural information gained by using cryoEM has allowed us to classify these anti-IC Abs into four distinct classes based on epitope, furthering our understanding of understudied Ab2ɑ anti-idiotypic antibodies (Fig. S11). Additional studies and the generation of anti-IC mAbs will be essential to further our understanding of how these antibodies are elicited by the immune system, engage with BCRs, and determine if these anti-IC Abs are autoreactive and may result in pathology.

While much remains to be understood about these anti-IC Abs, they do represent an additional non-neutralizing immune response that may compete with functional Ab responses. Emerging EMPEM data from human clinical trials employing SOSIP immunogens suggest that the phenomenon we describe herein will also be borne out in humans. Hence, our data provide a basis for vaccine redesign to potentially suppress or exploit the immune complex effect on shaping Ab responses.

## Acknowledgements

The authors thank Bill Anderson, Will Lessin, and Hannah Turner from Scripps Research for their help with EM experiments. We thank Lauren Holden for their help in preparing this article for submission and to Gabriel Ozorowski for reviewing this manuscript and for their help depositing data. We also thank Mitch Brinkkemper for managing serum sample shipments. Finally, the authors would like to thank Sean Mulligan from the Pacific Northwest Cryo-EM Center (PNCC) for his support in the acquisition of several cryoEMPEM datasets.

## Author Contributions

Conceptualization: AA, ABW, SB Methodology: AA, SB

Formal Analysis: SB, AA, DMG Investigation: SB, AA, LMS, PJMB

Visualization: SB, DMG

Funding Acquisition: ABW, RWS

Project Administration: AA, SB, ABW

Supervision: AA, ABW

Writing – Original Draft: SB, ABW

Writing – Review and Editing: SB, ABW, AA, RWS, LMS, PJMB, DMG

## Data and Materials Availability

All data associated with this study are presented in the paper or supplementary materials. Maps generated from the electron microscopy data are deposited in the Electron Microscopy Databank under the accession IDs EMD-43838, EMD-43968, EMD-43967, EMD-43999, EMD-43935, EMD-43981, EMD-43998, EMD-43894, EMD-43895, EMD-43896, EMD-43909, EMD-43910, EMD-43911, EMD-43915, EMD-43916, EMD-43917, EMD-43918, EMD-43919, EMD-44076. Atomic models corresponding to these maps have been deposited in the Protein Data Bank under the accession IDs 9ATZ, 9AXK, 9AXI, 9AYV, 9AXD, 9AY6, 9AYS. The raw data reported in this study will be shared by the corresponding author upon request.

## Materials and Methods

### Expression and Purification of BG505-SOSIP, 16055-SOSIP, B41-SOSIP, and CH505/BG505-SOSIP Constructs

Constructs containing 16055 SOSIP.v8.3, B41 SOSIP.v4.1, CH505/BG505 SOSIP.v8.1, and BG505 SOSIP.v5.2 N241/N289 genes were codon optimized for mammalian cell expression and subcloned into a pPP14 vector and expressed in 293F cells (Thermo Fisher Science) using polyethylenimine. After incubation for six days, the transfected cells were harvested by centrifugation (7,000 RPM for 60 minutes at 4°C) and the supernatant was cleared by vacuum filtration (0.45 µm filtration units, Millipore Sigma). The expressed SOSIPs were purified as described previously (*13*, *25*) using PGT145 immunoaffinity chromatography (Sepharose 4B resin, GE Healthcare Life Sciences) and eluted with 3 M MgCl_2_. Eluate was collected with an equal volume of TBS buffer. Affinity purified protein was concentrated and buffer exchanged into TBS (100 kDa ultrafiltration units, Amicon) and subjected to size-exclusion chromatography (SEC) on a HiLoad 16/600 Superdex 200 pg column (Cytiva). Samples were concentrated to 1 mg/mL and stored at 4°C.

### Immunization Experiments

The rabbit Clade C 16055 SOSIP.v8.3 and Clade C/Clade A CH505/BG505 SOSIP.v8.1 immunizations and blood draws were performed at Covance Research Products Inc. (Denver, PA, USA). Immunizations were performed under compliance of all relevant ethical regulations for animal testing and research. The study received ethical approval from the Covance Institutional Animal Care and Use Committee with approval number C0038-19. The female New Zealand White rabbits referenced in this study were immunized intramuscularly at weeks 0, 4, and 20 with 30 μg of 16055 SOSIP.v8.3 or CH505/BG505 SOSIP.v8.1 trimer formulated with Adjuplex adjuvant (Empirion). Blood draws were performed on the day of immunization and at weeks 2, 6, 12, and 22. For additional immunization details, see Brouwer et al., 2021 (*11*).

The rabbit Clade B B41 SOSIP.v4.1 immunizations and blood draws were performed at Covance Research Products Inc. (Denver, PA, USA). Immunizations were performed under compliance of all relevant ethical regulations for animal testing and research. The study received ethical approval from the Covance Institutional Animal Care and Use Committee with approval number C0045-15. The female New Zealand White rabbit referenced in this study was immunized intramuscularly at weeks 0, 4, and 20 with 22 μg of B41 SOSIP.v4.1 formulated with Iscomatrix^TM^ adjuvant (CSL Biotherapies, Inc). Blood draws were performed on the day of immunization and at weeks 6, 8, 10, 12, 16, and 22. For additional immunization details, see Torrents de la Peña et al., 2018 (*12*).

The rhesus macaque Clade A BG505 SOSIP.v5.2 N241/N289 immunizations and blood draws were performed at the Yerkes National Primate Research Center, Atlanta, GA, USA. All procedures were approved by Emory University Institutional Animal Care and Use Committee protocol 201700723, and animal care facilities are accredited by the U.S. Department of Agriculture and the Association for Assessment and Accreditation of Laboratory Animal Care International. The rhesus macaque referenced in this study was immunized at weeks 0, 8, 24, and 36 with BG505 SOSIP.v5.2 N241/N289 (100 μg per dose). Immunogens were formulated with Matrix-M^TM^ (Novavax, Inc.; 75 µg per dose) for the first three immunizations (weeks 0, 8, and 24). For the week 36 immunization, the adjuvant was changed to SMNP (Darrell Irvine Lab, MIT; 750 U per dose). Blood draws were performed biweekly. For additional immunization details, see Antanasijevic et al., 2021 (*13*).

### Isolation of IgG and Fab Preparation for nsEMPEM

IgGs were purified from ∼2 mL of plasma with an equal volume of Protein A Sepharose resin (GE Healthcare Life Sciences) and incubated with the resin for three days at 4°C. The resin was washed three times with 10 column volumes (CV) of PBS and eluted with 5 CV 0.1 M glycine pH 2.5 into a tube containing 1 M Tris-Cl pH 8.0. Samples were filtered with 0.22 µm filtration units, (Steriflip, Millipore Sigma) concentrated to ∼4-6 mg/mL and buffer exchanged into PBS pH 7.4 (30 kDa ultrafiltration units, Amicon).

The desired amount of IgG was added to the digestion buffer (100 mM Tris + 10 mM EDTA + 10 mM cysteine, pH 7.4) and digested for 5 h at 37°C using papain (1 mg/mL final concentration). Digestion was quenched using iodoacetamide. The digested Fabs were buffer exchanged into TBS (10 kDa ultrafiltration units, Amicon). Fabs were purified using SEC on a Superdex 200 Increase 10/300 GL column (Cytiva).

Fab complexes were assembled using 1 mg of purified polyclonal Fab and 20 µg of the corresponding SOSIP antigen from immunization (BG505 SOSIP.v5.2 N241/N289, 16055 SOSIP.v8.3, B41 SOSIP.v4.1, CH505/BG505 SOSIP.v8.1). The reactions were incubated ∼18 hours at room temperature. Immune complexes were purified from remaining Fab using SEC (Superose 6 Increase Column, Cytiva). Fractions were concentrated (10 kDa ultrafiltration units, Amicon) and immediately used for negative stain EM as described below.

### Negative Stain Electron Microscopy (nsEM)

Negative stain experiments were performed as described previously (*5*, *13*). Purified complexes were diluted to ∼40-80 µg/mL and applied to carbon-coated 400-mesh copper grids that were glow-discharged at 15 mA for 25 sec. Sample was applied for 15 sec and blotted. Grids were negatively stained using 2% w/v uranyl-formate for 40 sec and blotted. Data was collected on a Tecnai TF20 electron microscope operating at 200 keV with a pixel size of 1.77 Å. Micrographs were recorded using a Tietz 4k x 4k TemCam-F416 CMOS detector. The Leginon automated imaging interface was used for data acquisition (*26*) and initial data processing was performed using the Appion data processing suite (*27*). Relion/3.0 was used for 2D and 3D classification steps (*28*).

### nsEMPEM Data Processing

Approximately 100,000 particles were auto-picked using the Appion data processing package. These particles were extracted and underwent 2D classification using Relion/3.0 into 200 classes (25 iterations). Particles with antigen-Fab qualities were selected for 3D analysis. The initial round of 3D classification was performed using 40 classes. Particles from similar looking classes were combined and reclassified, these subgroups with unique structural features were further processed using 3D auto-refinement. UCSF Chimera 1.15 (*14*) was used to visualize and segment 3D refined maps for figures.

### Preparation of Fab and Complexes for CryoEMPEM

Polyclonal Fab samples from individual immunizations were complexed with 200 µg of the corresponding SOSIP antigen (BG505 SOSIP.v5.2 N241/N289, 16055 SOSIP.v8.3, B41 SOSIP.v4.1, CH505/BG505 SOSIP.v8.1) and incubated for ∼18 hours at room temperature. Complexes were purified using SEC (HiLoad 16/600 Superdex 200 pg Column, Cytiva) and concentrated to 5-7 mg/mL (10 kDa ultrafiltration units, Amicon) for application onto CryoEM grids.

### CryoEMPEM Grid Preparation and Imaging

Grids were prepared using a Vitrobot Mark IV (Thermo Fisher Scientific). The chamber temperature was set to 10 °C with 100% humidity and variable blot times between 4-7 sec. Blot force was set to zero and wait time was set to 10 sec. Two types of grids, UltAuFoil R. 1.2/1.3 (Au, 300-mesh; Quantifoil Micro Tools GmbH) and Quantifoil R 2/1 (Cu, 400-mesh; Quantifoil Micro Tools GmbH) were used for freezing. Before sample application, grids were treated with Ar/O2 plasma (Solarus plasma cleaner, Gatan) for 10 sec.

Lauryl maltose neopentyl glycol (LMNG) at a final concentration of 0.005 mM was mixed with the sample and 3 µL of the solution was immediately loaded onto the grid. After blotting, the grids were plunge frozen in liquid nitrogen-cooled liquid ethane. Grids were loaded into an FEI Titan Krios electron microscope (Thermo Fisher Scientific) operating at 300 kV. The microscope is equipped with a K2 Summit direct electron detector (Gatan) and sample autoloader. Leginon software was used for automated data collection (*26*). Specific details on imaging and data collection for individual polyclonal complexes can be found in Supplementary Table 1.

### CryoEMPEM Data Processing

MotionCor2 (*29*) was used to align and dose-weight micrographs. Aligned micrographs were uploaded to cryoSPARCv2.15 (*30*) where initial data processing was performed. GCTF (*31*) was used to estimate the CTF parameters. Immune complexes were picked using the template picker, extracted, and underwent three rounds of 2D classification to remove bad particle picks. A selected subset of particles were transferred to Relion/3.0 (*28*) for additional data processing.

After a round of 2D classification in Relion, the particles belonging to classes with the trimer-Fab immune complexes were selected and subjected to 3D refinement (C3 symmetry, soft solvent mask around the trimer). A low pass filtered map of BG505 SOSIP trimer was used as an initial model for all 3D steps to avoid initial model bias during reconstruction. The aligned particles were symmetry expanded around the C3 axis to collapse all epitope-paratope interfaces onto a single protomer, simplifying 3D classification and assuring all possible epitopes are considered.

To prevent symmetry-related copies of individual particles from aligning to one another, particle alignment was constrained in the following 3D classification and refinement steps. 3D classification steps were performed without image alignment (--skip_align, T=16) and 3D refinement steps were performed with local angular searches only (3.7° per iteration).

The first round of 3D classification utilized a sphere mask with a diameter of 80 Å positioned around the epitope-paratope interface of interest containing density that corresponds to a bound polyclonal Fab. This is performed separately for each epitope area of interest where polyclonal Fab density can be detected. This sphere mask allows for sorting particles based on the absence or presence of Fabs at each location on the antigen as well as separating unique polyclonal Fabs with different orientations and epitope footprints. The number of 3D classes was adjusted for each epitope depending on the relative occupancy of the bound polyclonal Fabs (10-30 classes). Classes of particles with structurally unique polyclonal fabs were selected separately and processed independently for the following steps.

All of the selected particle subsets underwent a round of 3D refinement with a soft-solvent mask around the specific trimer-Fab complexes. A second round of 3D classification was performed with highest quality and resolution particles from refinement. A mask of the polyclonal Fab was used to search for classes of particles with highest resolution and best reconstruction of the individual Fab (T=16; 5-10 classes). A second round of 3D refinement was performed on the selected particles using the same mask as previous refinements.

A third round of 3D classification was used to refine the resolution of the epitope-paratope interface using either the individual Fab mask used during the last refinement or using an 80 Å sphere mask targeting the epitope-paratope interface (T=16; 3-5 classes). The highest resolution class was selected for each unique complex and 3D refined according to previous rounds. A final round of 3D classification was performed to select a final set of particles with the highest resolution and quality in the trimer, epitope-paratope interface, and Fab. This was performed with a mask of the entire trimer-Fab complex (T=16; 3 classes). The best class was selected, 3D refined, and postprocessed using a solvent mask around the complex (correction for the modulation transfer function (MTF) parameters was performed). The postprocessed maps were used for model building and submission to the EMDB. Additional data processing information is presented in Supplementary Table 1.

### CryoEMPEM Model Building and Refinement

Postprocessed maps from Relion were used for model building and refinement. An initial model of 16055 NFL trimer (PBD ID: 6P65) was used to build the antigen-corresponding parts of the 16055 SOSIP.v8.3 anti-idiotypic antibody structure. The sequence was adjusted to match the exact 16055 SOSIP.v8.3 immunogen used in the experiment. Subsequent 16055 SOSIP.v8.3 structures (N625+Base, N611+Base) used the 16055 SOSIP.v8.3 anti-idiotypic antibody structure as an initial model. An initial model of B41 SOSIP.v664 (PDB ID: 6OKP) was used to build the antigen-corresponding parts of the B41 SOSIP.v4.1 anti-idiotypic antibody structure. The sequence was altered to include the appropriate SOSIP.v4.1 mutations.

Due to the lower resolution of the anti-idiotypic antibody epitope/paratope interface, models of the individual interface and V5 Fabs were separately built and combined to create an initial model for the Rh.33203 Class IV anti-idiotypic antibody structure. An initial model of BG505 SOSIP.v5.2 N241/N289 (PDB ID: 7L8X) was used to build the antigen-corresponding portion of the BG505 SOSIP.v5.2 N241/N289 interface antibody structure. This model was used as the initial model for the BG505 SOSIP.v5.2 N241/N289 V5 antibody. The BG505 SOSIP.v5.2 N241/N289 interface antibody model was combined with the V5 Fab and used as an initial model for the BG505 SOSIP.v5.2 N241/N289 anti-idiotypic antibody structure.

The CH505/BG505 SOSIP.v8.1 chimera used in this study lacked a corresponding model available in the PBD. The sequence of this CH505/BG505 chimera was used as a query for NCBI BLASTp (*32*). A model of BG505 SOSIP.v5.2(7S) (PBD ID: 7MEP) was used as the initial model and relaxed into the reconstructed cryoEMEPM map. The sequence was mutated to match the CH505/BG505 SOSIP.v8.1 immunogen sequence used in the immunization.

Initial models of Fabs were generated from PDB ID: 4TZO. All amino acids were mutated to alanine and docked into each SOSIP map in UCSF Chimera to generate a starting model for refinement. Antibody heavy (H) and light (L) chains were assigned by comparing the CDR2 and CDR3 lengths and conformations of framework regions between the heavy and light chains in the reconstructed CryoEM maps. Iterative rounds of manual model refinement in Coot (*33*) and automated model refinement in Rosetta (*34*, *35*) were used to refine models into the CryoEM maps. CDR length was altered by inserting or deleting alanine residues and refining the model to match the corresponding CryoEM map. Final models were evaluated using MolProbity (*36*) and EMRinger (*37*), refinement statistics are shown in Supplementary Table 1. The structures were submitted to the PDB (see table S2 for EMDB and PDB accession codes).

### B Cell Repertoire Sequencing of 16055 2464

Peripheral blood mononuclear cells (PBMCs) were harvested from New Zealand White rabbits immunized with 16055 SOSIP.v8.3 at W21 (*11*). We were able to obtain approximately 10 million PBMCs from 16055 2464 cryopreserved in 45% FBS, 45% RPMI, and 10% DMSO. These PBMCs were sent to Abterra Biosciences (San Diego, CA) for bulk B cell receptor (BCR) sequencing using their Reptor™next generation sequencing (NGS) platform. Briefly, RNA was extracted and used to synthesize cDNA. IgG was amplified and the full variable region was sequenced. Data was filtered and error correction was performed. Unfortunately, the PBMCs had undergone extensive RNA degradation at some point between collection and amplification which limited the number of viable sequences. Only 518 heavy chain and 36,242 light chain sequences were generated.

### Automated Model Building Using ModelAngelo

ModelAngelo is a tool developed for automated atomic model building in cryoEM maps (*15*). Using a machine learning approach, it combines information from the cryoEM map with information from about protein structure and sequence to build an accurate model. ModelAngelo can also be used to build an atomic model into areas of unknown sequences, which is particularly useful for pAbs which lack sequence information. The program was installed from GitHub and used to generate a model for the 16055 2464 anti-IC and V2 pAbs (3.4 Å resolution map). The output model and .hmm profiles were used to search the unpaired BCR sequence database using HMMER (*38*). The top scoring hits were analyzed in Coot (*33*). Sequences for the anti-IC heavy and light chains and V2 heavy chain that closely matched the EM map were isolated and refined to fit the cryoEM map. Unfortunately, no probable V2 light chain sequence was identified in the repertoire.

### Frequency Analysis

To determine the amino acid frequency for the 16055 2464 anti-IC and V2 pAbs, a multiple sequence alignment using Clustal Omega 1.2.4 (*39*, *40*) was generated using IgK and IgH sequence databases from B cell repertoire sequencing. We calculated the amino acid probability distribution 𝑃(𝑋_𝑖_|𝐴) to identify highly conserved regions where 𝑚 is the number of sequences, 𝑖 is the alignment position, and 𝐴 is an amino acid.

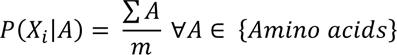

To identify sequence motifs based on amino acid properties, we grouped amino acids with similar properties into four groups (𝐺): polar (*STNQ*), charged (*RHKDE*), hydrophobic (*AVILMFYW*), and aromatic (*FYWH*). For each group, the probability distribution per alignment position was calculated where 𝑚 is the number of sequences, 𝑖 is the alignment position, and 𝐺 is one of the four groups.

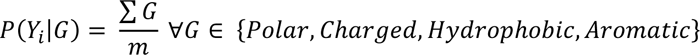

