## Supplemental Material for "Anti-Immune Complex Antibodies are Elicited During Repeated Immunization with HIV Env Immunogens"

**Fig. S1.**

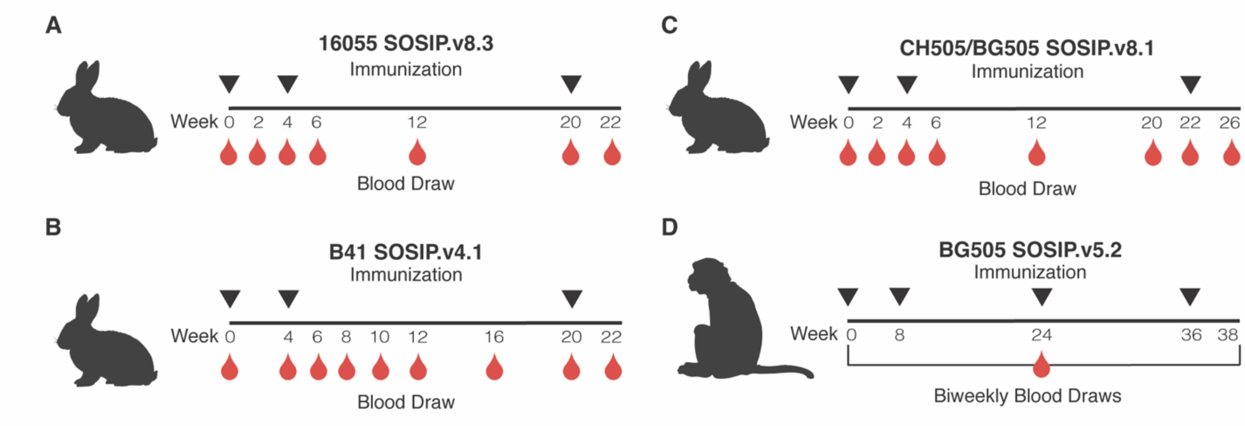


**Supplemental Figure 1. Immunization schedules.** Rabbits were immunized three times with: **A)** 16055 SOSIP.v8.3, **B)** CH505/BG505 SOSIP.v8.1, or **C)** B41 SOSIP.v4.1. Blood was drawn at a series of time points post-immunization. **D)** Rhesus macaques were immunized four times with BG505 SOSIP.v5.2 N241/N289 and blood draws were performed biweekly. See the methods for additional immunization details.

**Fig. S2**

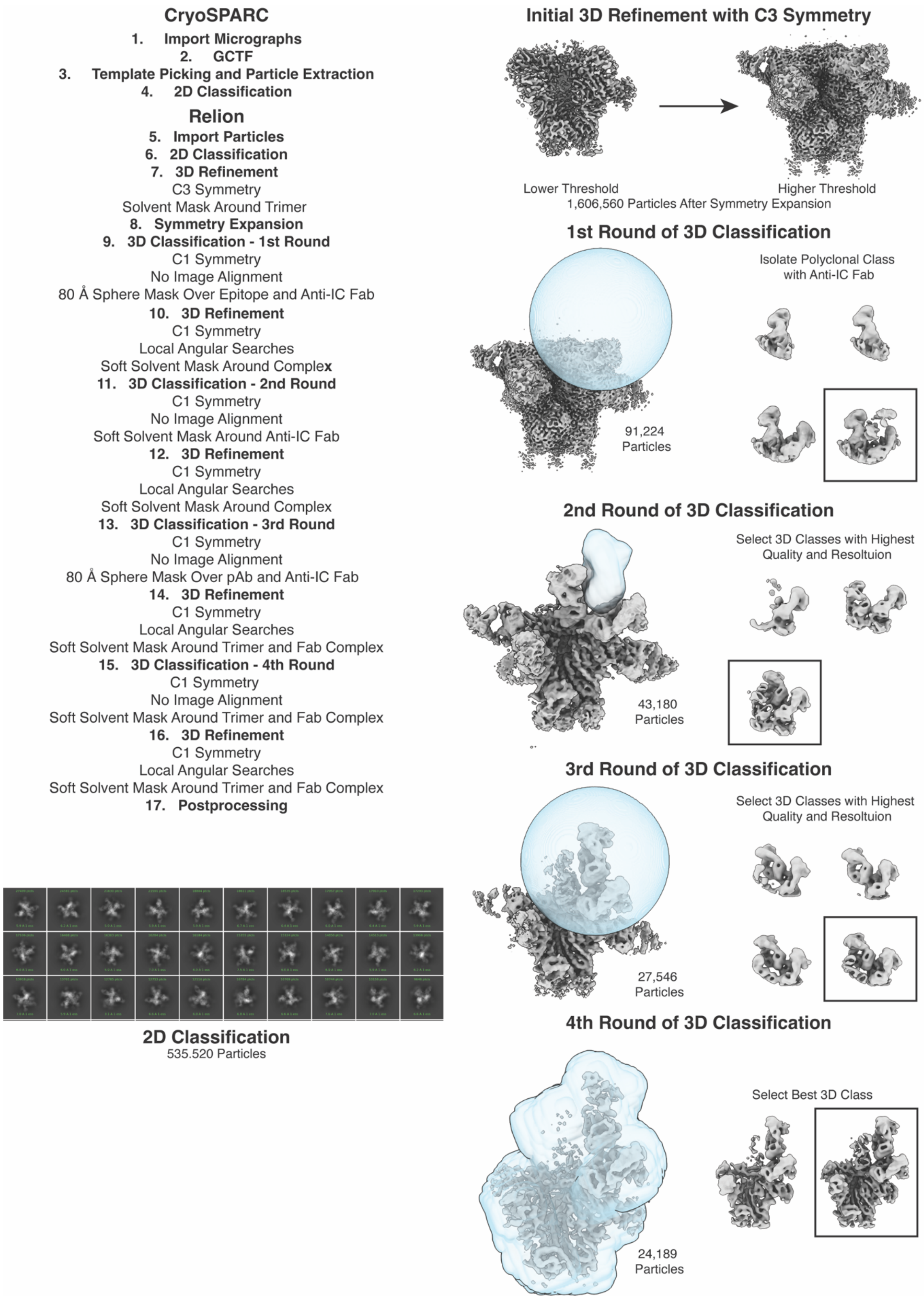
CryoSPARC
1. Import Micrographs
2. GCTF
3. Template Picking and Particle Extraction
4. 2D Classification
Relion
5. Import Particles
6. 2D Classification
7. 3D Refinement
C3 Symmetry
Solvent Mask Around Trimer
8. Symmetry Expansion
9. 3D Classification - 1st Round
C1 Symmetry
No Image Alignment
80 Å Sphere Mask Over Epitope and Anti-IC Fab
10. 3D Refinement
C1 Symmetry
Local Angular Searches
Soft Solvent Mask Around Complex
11. 3D Classification - 2nd Round
C1 Symmetry
No Image Alignment
Soft Solvent Mask Around Anti-IC Fab
12. 3D Refinement
C1 Symmetry
Local Angular Searches
Soft Solvent Mask Around Complex
13. 3D Classification - 3rd Round
C1 Symmetry
No Image Alignment
80 Å Sphere Mask Over pAb and Anti-IC Fab
14. 3D Refinement
C1 Symmetry
Local Angular Searches
Soft Solvent Mask Around Trimer and Fab Complex
15. 3D Classification - 4th Round
C1 Symmetry
No Image Alignment
Soft Solvent Mask Around Trimer and Fab Complex
16. 3D Refinement
C1 Symmetry
Local Angular Searches
Soft Solvent Mask Around Trimer and Fab Complex
17. Postprocessing
2D Classification
535.520 Particles
Initial 3D Refinement with C3 Symmetry
Lower Threshold
Higher Threshold
1,606,560 Particles After Symmetry Expansion
1st Round of 3D Classification
Isolate Polyclonal Class with Anti-IC Fab
91,224 Particles
2nd Round of 3D Classification
Select 3D Classes with Highest Quality and Resoltuion
43,180 Particles
3rd Round of 3D Classification
Select 3D Classes with Highest Quality and Resoltuion
27,546 Particles
4th Round of 3D Classification
Select Best 3D Class
24,189 Particles

**Supplemental Figure 2. Data processing workflow and masking strategy.** Representative data processing overview schematic for EMD-43999 performed in CryoSPARC 2.15 (*30*) and Relion 3.0 (*28*). Similar strategies were employed for other maps produced in this work. Representative 2D classes are shown on the bottom left taken from EMD-43999. The masking strategy employed for data processing is shown on the right with EMD-43999 used as an example. The first round of 3D classification uses an 80 Å sphere mask over the epitope-paratope interface to isolate particles containing the anti-immune complex antibody. A second round of 3D classification using a Fab mask of the anti-immune complex antibody is used to select classes of high-quality particles. A third round of 3D classification using an 80 Å sphere mask is used to select the highest resolution class of particles. A final round of 3D classification is used to select particles with high resolution in the trimer, epitope-paratope region, and Fabs.

Fig. S3.

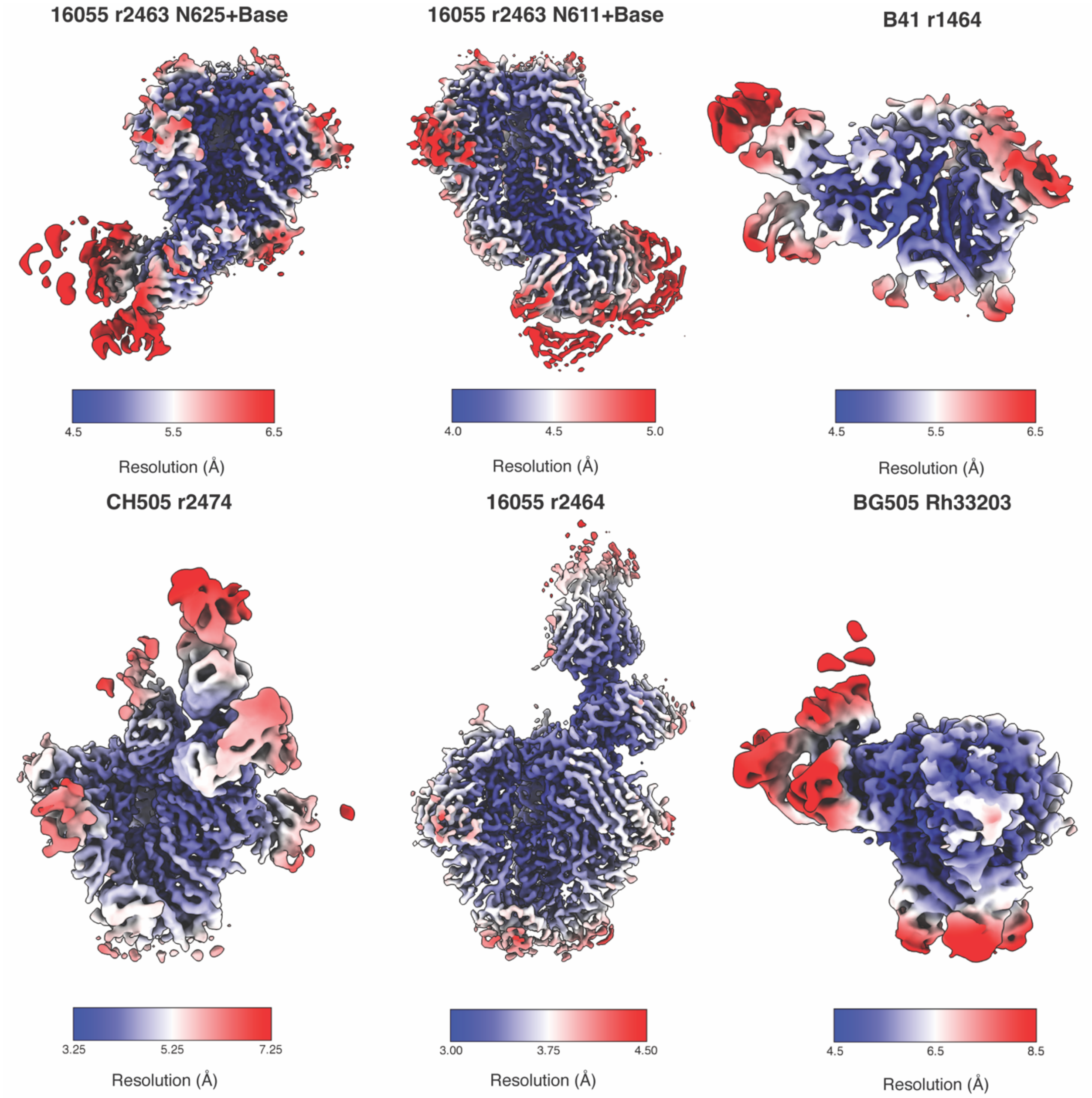


**Supplemental Figure 3. Local resolution plots for EM maps of anti-immune complex antibodies.** Local resolution was calculated according to a 0.143 FSC threshold in Relion 3.0 (*28*) and visualized in ChimeraX (*14*).

**Fig. S4.**

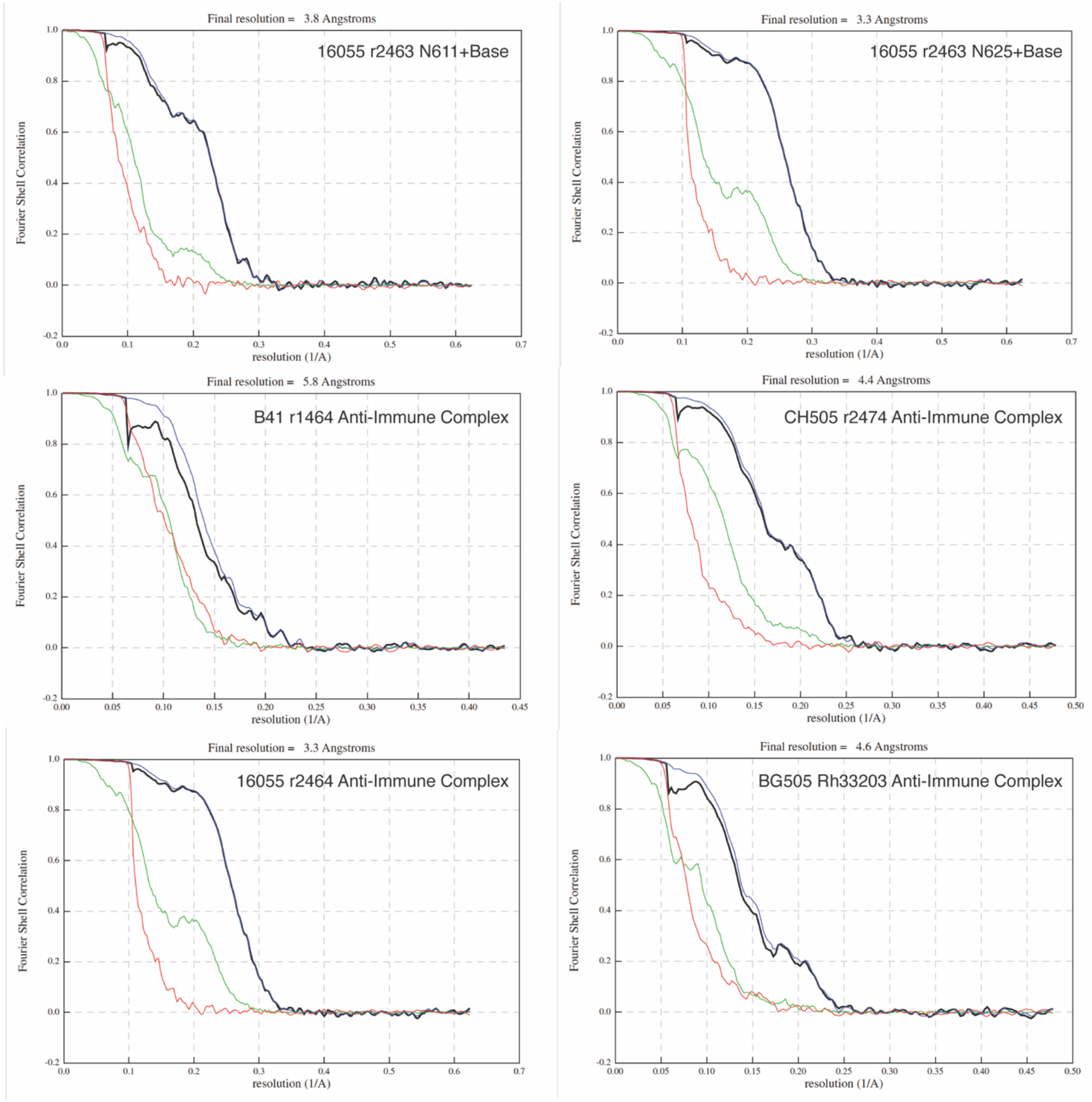


**Supplemental Figure 4. FSC plots for EM maps of anti-immune complex antibodies.** Relion 3.0 (*28*) was used to generate FSC plots. Reported resolutions coincide with a 0.143 FSC cutoff. The red line represents the phase randomized masked map FSC, the green line represents the unmasked map FSC, the blue line represents the masked map correlation, and the black line represents the corrected FSC.

**Fig. S5.**

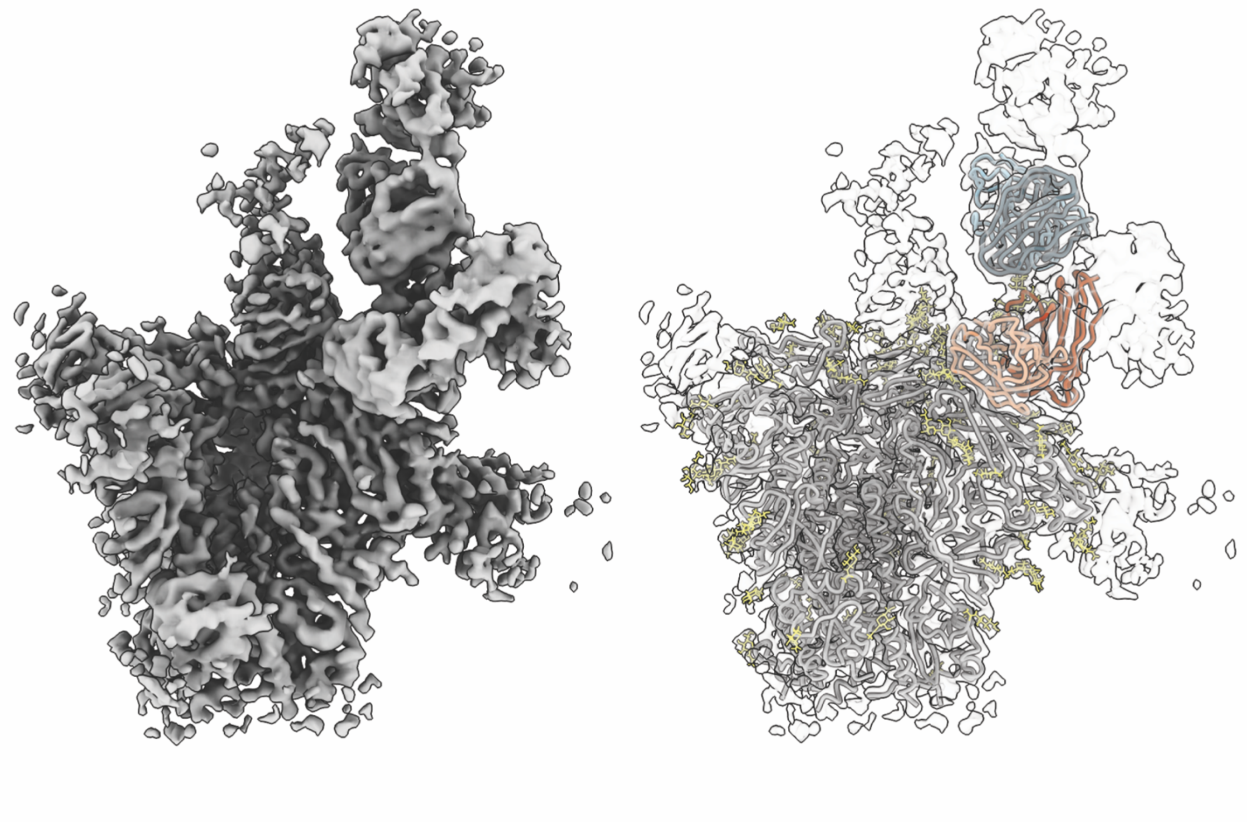

**Supplemental Figure 5. The Class II CH505/BG505 r2474 map contains an additional antibody bound to the V2 apex epitope.** During cryoEMPEM data processing for CH505/BG505 r2474 a number of classes which contained an additional pAb bound to the V2 apex epitope were observed. A majority of the particles containing the anti-immune complex (anti-IC) antibody also contained this V2 apex pAb. The final map (left) shows diffuse low-resolution density for this additional pAb which does not appear to contact the V1/V3 pAb (orange) or the anti-IC antibody (blue) and therefore was not modeled in the final structure (right).

**Fig. S6.**

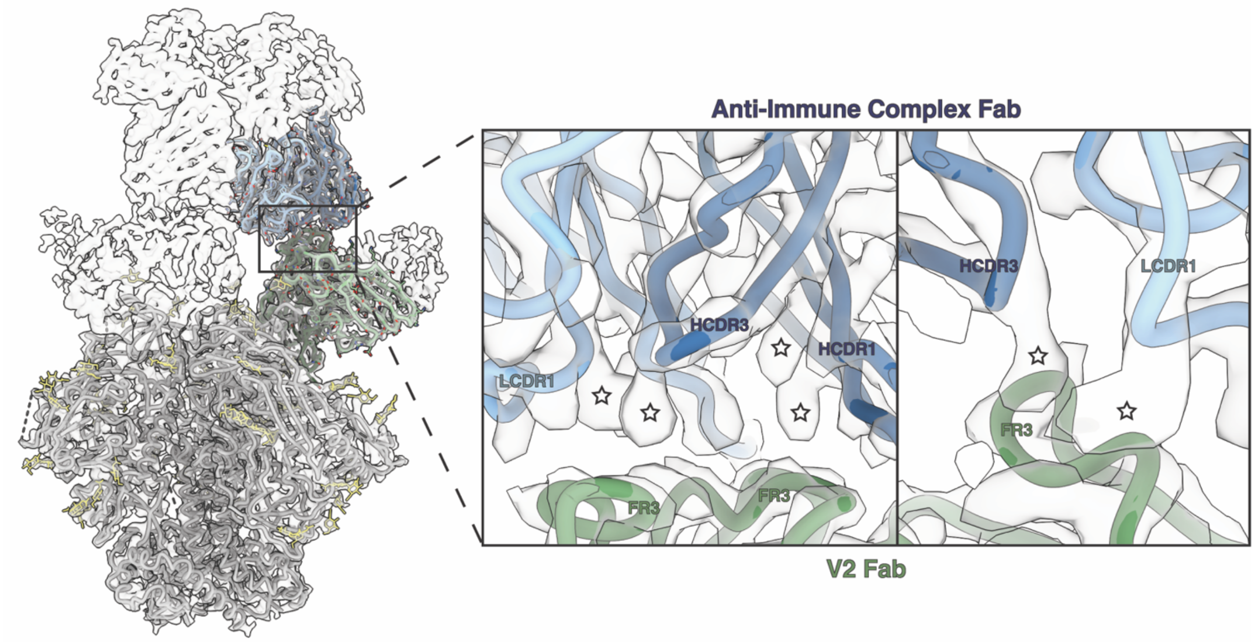


**Supplemental Figure 6. The epitope-paratope interface of the Class III 16055 r2464 V2 pAb and anti-immune complex antibody contains aromatic amino acids.** The anti-IC Ab (blue) contacts the V2 pAb (green) epitope using a paratope dominated by side chain densities that correspond to aromatic amino acids. These side chain densities are marked with a star. LCDR1 (light blue) contains side chain density that resembles a tryptophan residue. HCDR3 and HCDR1 (dark blue) contain a number of aromatic side chains that appear to be tyrosine or phenylalanine residues.

**Fig. S7.**

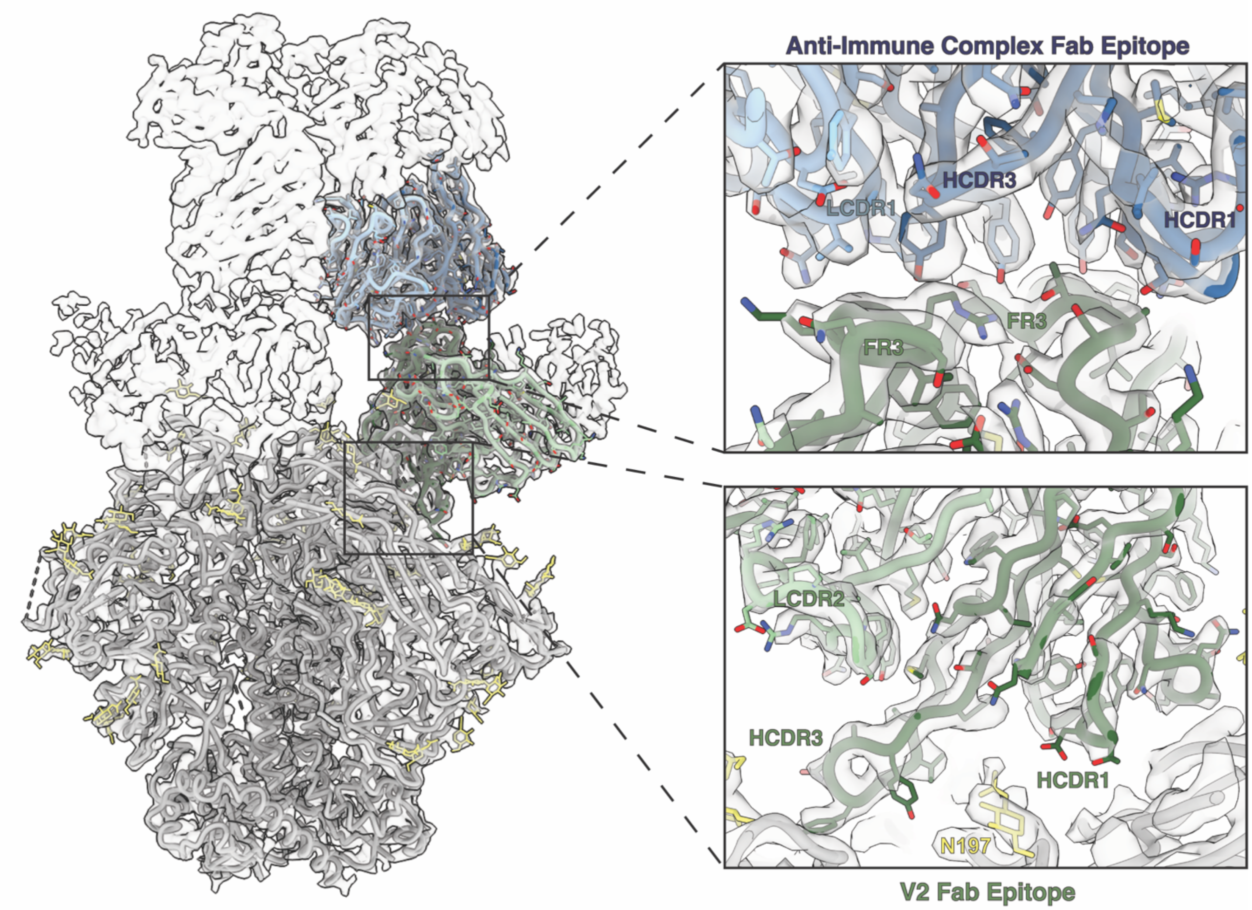


**Supplemental Figure 7. Predicted sequences of Class III 16055 r2464 V2 and anti-immune complex Fabs.** ModelAngelo (*15*) was used to generate predicted sequences for the Class III anti-immune complex antibody and the V2 pAb to which it binds. The output sequence produced by ModelAngelo was used to search a NGS database of unpaired BCR sequences from this animal. The top sequences were aligned in Clustal Omega (*39,40*) and used to generate a consensus sequence based on features in the EM map. Heavy (dark blue) and light chain (light blue) sequences were identified for the anti-immune complex antibody and modeled into the map above. Additionally, a heavy chain sequence for the V2 pAb was identified and modeled (dark green). However, the light chain V2 pAb NGS search did not return viable hits. The ModelAngelo output sequence is thus modeled in its place (light green). The paratope of the anti-immune complex antibody is enriched for aromatic amino acids (see top right inset), many of which contact framework regions of the V2 Fab.

**Fig. S8.**

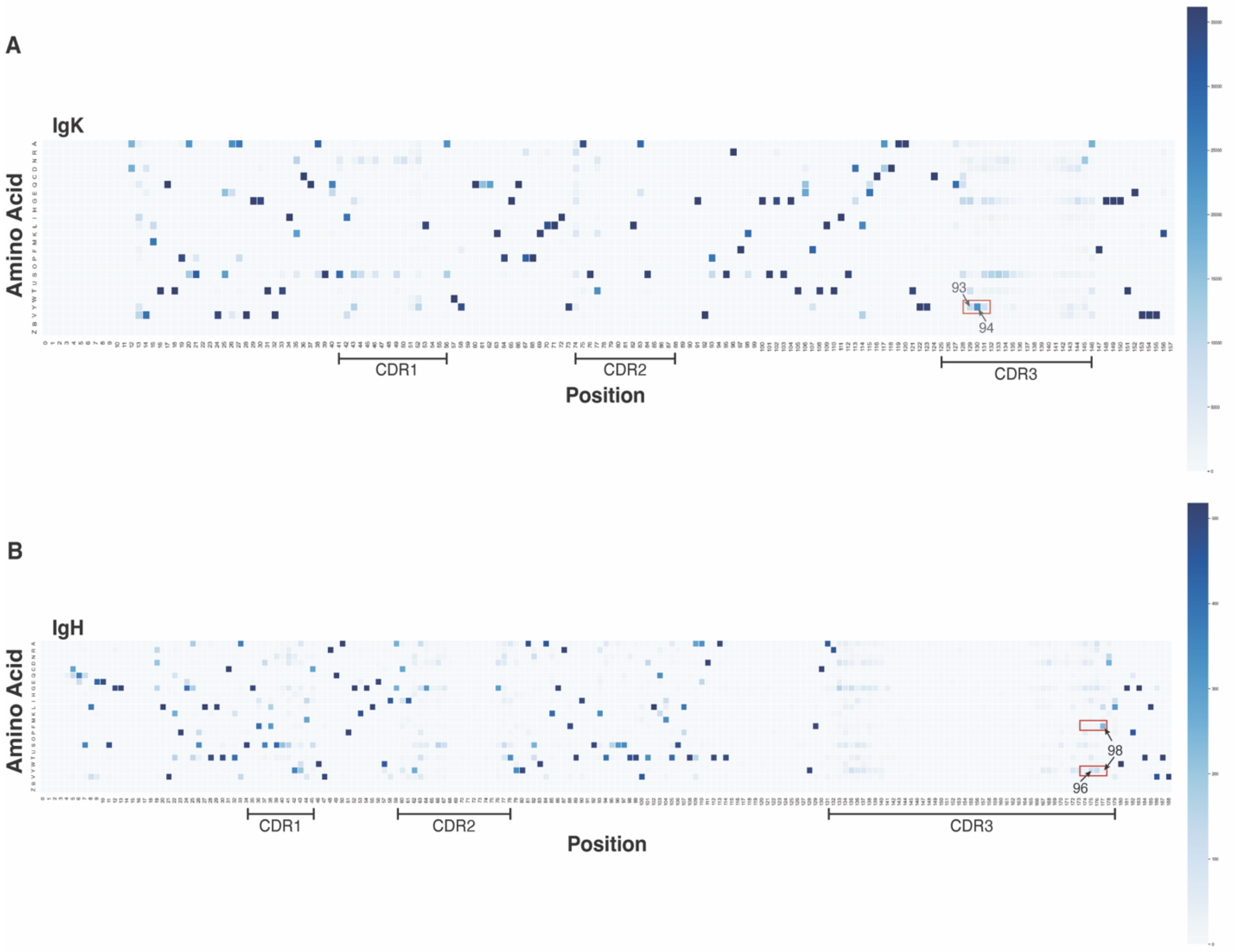


**Supplemental Figure 8. Amino acid probability distribution for the Class III 16055 r2464 anti-immune complex antibody**. Unpaired B cell repertoire (BCR) sequencing was performed on PBMCs obtained from 16055 r2464 and used to determine the frequency of aromatic amino acids in the anti-immune complex antibody isolated from this animal. The amino acid probability distribution was calculated relative to the number of heavy and light chain repertoire sequences, with darker shades of blue representing a higher frequency of that amino acid appearing at any given position. Position numbering is based on the multiple sequence alignment and does not correspond to the residue number in the 16055 r2464 model. For calculation details, see the methods. **A)** The light chain CDR3 contains two aromatic residues at positions 93 and 94 (boxed in red). The frequency of an aromatic amino acid (Tyr, Phe, Trp, His) occurring at these positions are 26% and 70% respectively. There is no significant frequency for Phe at these positions, indicating the residues are likely Tyr. **B)** The heavy chain CDR3 also contains two aromatic residues at positions 96 and 98. The frequency of an aromatic amino acid occurring at these positions are 25% and 55% respectively. The most probable identity of residue 96 is Tyr (lower red box) and for residue 98 is Phe (top red box).

**Fig. S9.**

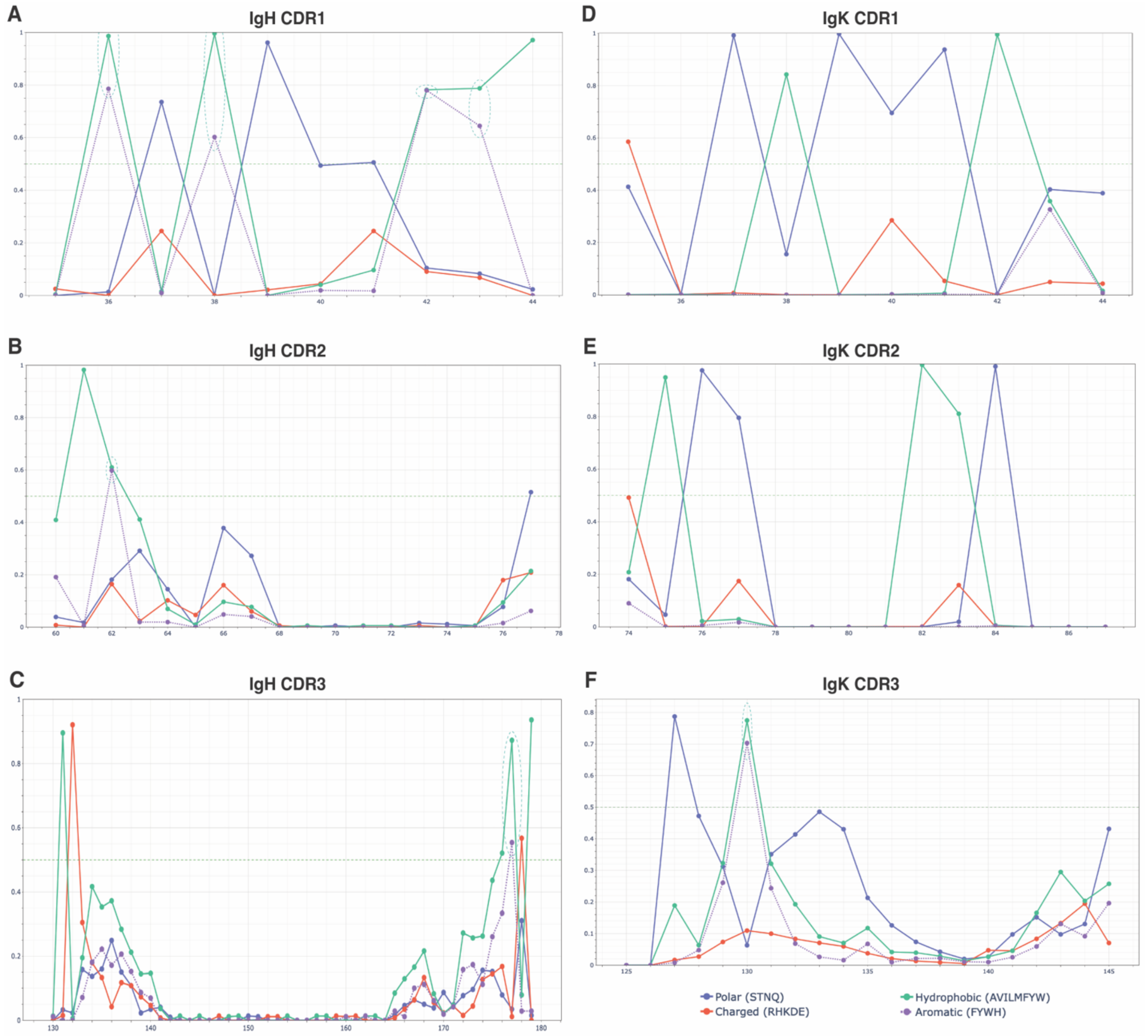


**Supplemental Figure 9. CDR sequence motifs for the Class III 16055 r2464 anti-immune complex antibody**. Using the BCR sequencing data from 16055 r2464 we were also able to determine sequence motifs based on amino acid properties (polarity, charge, hydrophobicity, aromaticity) in the CDR regions of the anti-IC Ab. Positions where aromatic residues are above the probability threshold (0.5) are circled in green. Position numbering is based on the multiple sequence alignment and does not correspond to the residue number in the 16055 r2464 model. A) HCDR1 contains a number of probable aromatic amino acids at positions 36, 38, 42, and 43. The predicted sequence shows two aromatic residues, Phe36 and Tyr42, but no aromatic residues at Leu38 and Asp43. B) HCDR2 contains an aromatic amino acid at position 62 which is a Tyr in the predicted sequence. C) The sequence motif for HCDR3 shows one aromatic residue at position 177. The predicted sequence shows two aromatics, Tyr175 and Tyr177. The sequence motifs for LCDR1 (D) and LCDR2 (E) do not show significant likelihood of aromatic amino acids. F) The sequence motif for LCDR3 contains one aromatic residue at position 130. In the predicted sequence there are two aromatic residues, Tyr129 and Tyr130.

**Fig. S10.**

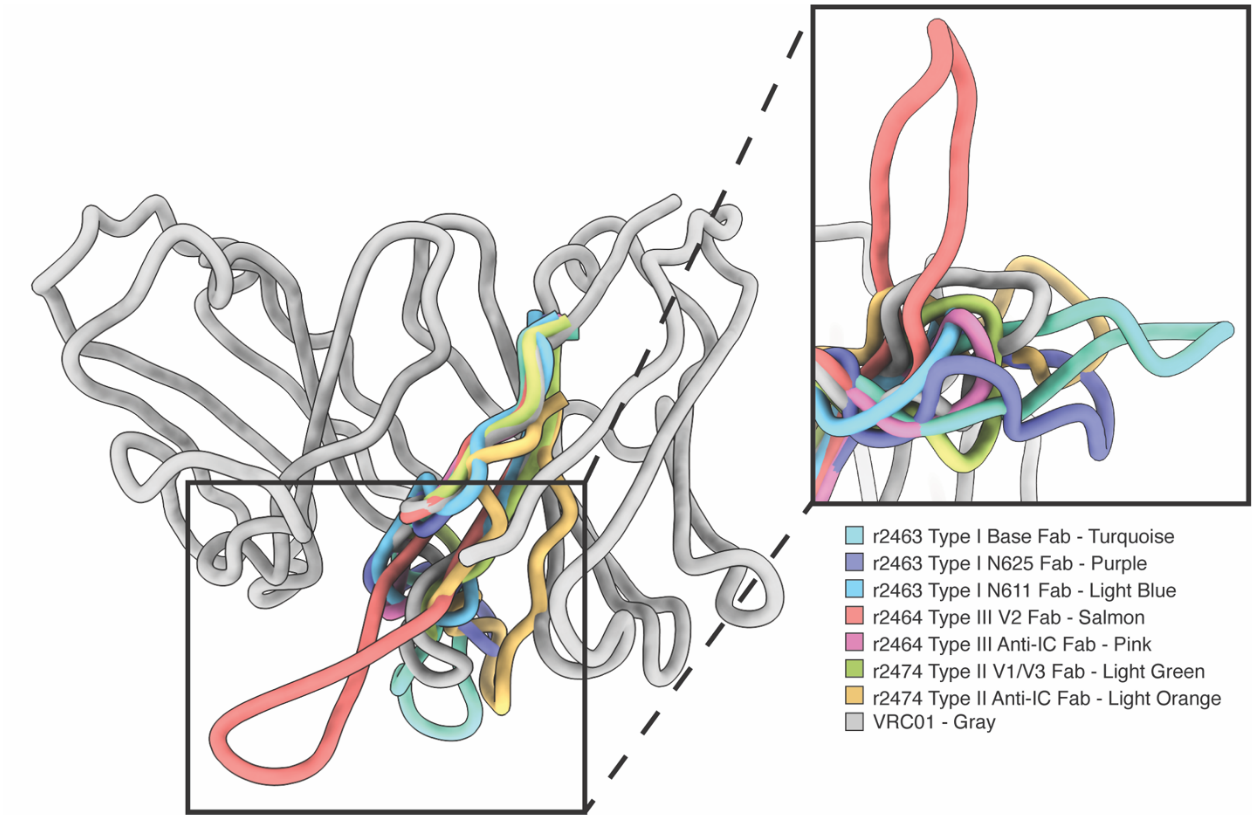


**Supplemental Figure 10. Anti-immune complex antibodies have shorter HCDR3 loops.** High local resolution in the epitope-paratope interface between the anti-immune complex antibody and pAbs bound to Env facilitated the determination of key anchor residues. These anchor residues were used to estimate HCDR3 length for Class I 16055 r2463, Class II CH505/BG505 r2474, and Class III 16055 r2464 models. HCDR3 loops are shown aligned to a model of VRC01 (PDB ID: 6MYY). The average length of Class II and Class III anti-immune complex antibodies shown here was 12.5 amino acids (n=2). The average length of antibodies bound to Env and Class I anti-immune complex antibodies (which have an epitope only composed of Env) was 17.4 amino acids (n=5).

**Fig. S11.**

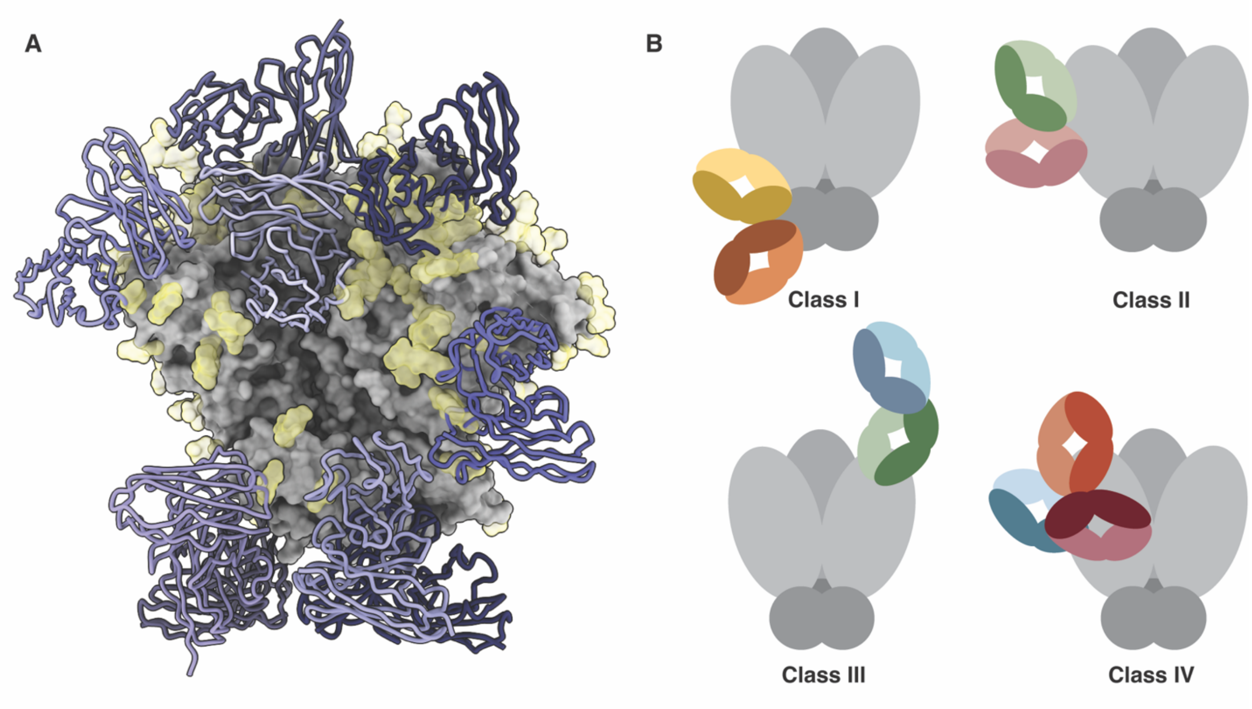


**Supplemental Figure 11. Epitope masking by immunodominant primary antibodies and heavy glycosylation prime secondary antibody responses which may elicit anti-immune complex antibodies.** HIV Env is heavily glycosylated which can mask neutralizing epitopes on the antigen surface. Immunodominant antibodies elicited after initial priming and boosting immunizations can mask the underlying peptide surface, directing secondary antibody responses to subdominant epitopes. A) N-linked glycans (yellow) and primary antibodies (purple) identified during cryoEMPEM are shown on one face of the HIV Env trimer. The available peptide surface is limited due to epitope masking, which primes secondary antibody responses to target subdominant epitopes that include N-linked glycans and framework regions of primary antibodies. B) There are four distinct classes of anti-immune complex antibodies that are elicited against primary antibodies. Class I anti-immune complex antibodies have an epitope composed of antigen but make non-specific contacts with one another along framework regions. Class II anti-immune complex antibodies have a neoepitope composed of both antigen and antibody framework regions. Class III anti-immune complex antibodies are canonical Ab2α anti-idiotypic antibodies and have an epitope composed entirely of antibody framework regions. Class IV anti-immune complex antibodies bind to an idiotope composed of framework regions from two separate antibodies.

**Table S1.**

**Supplemental Table 1. CryoEM data collection and model refinement statistics.**

| | **16055 SOSIP.v8.3 r2463 N611 + Base** | **16055 SOSIP.v8.3 r2463 N625 + Base** | **16055 SOSIP.v8.3 r2464** | **CH505/BG505 SOSIP.v8.1 r2474** | **BG505 SOSIP.v5.2 N241/N289 Rh33203 Full Complex** | **BG505 SOSIP.v5.2 N241/N289 Rh33203 Interface Fab** | **BG505 SOSIP.v5.2 N241/N289 Rh33203 V5 Fab** | **B41 SOSIP.v4.1 r1646** |
|---|---|---|---|---|---|---|---|---|
| **Access Codes** | | | | | | | | |
| PDB | 9AXK | 9AXI | 9ATZ | 9AYV | 9AYS | 9AXD | 9AY6 | - |
| EMDB | EMD-43968 | EMD-43967 | EMD-43838 | EMD-43999 | EMD-43998 | EMD-43935 | EMD-43981 | EMD-43894 |
| **Data Collection and Processing** | | | | | | | | |
| Microscope | TFS Titan Krios | TFS Titan Krios | TFS Titan Krios | TFS Titan Krios | TFS Titan Krios | TFS Titan Krios | TFS Titan Krios | FEI Talos Arctica |
| Detector | Gatan K3 | Gatan K3 | Gatan K2 | Gatan K2 | Gatan K2 | Gatan K2 | Gatan K2 | Gatan K2 |
| Nominal Magnification | 29,000 | 29,000 | 130,000 | 130,000 | 130,000 | 130,000 | 130,000 | 36,000 |
| Voltage (kV) | 300 | 300 | 300 | 300 | 300 | 300 | 300 | 200 |
| Electron Exposure ($e^-/Å^2$) | 40.3 | 40.3 | 50.0 | 49.8 | 49.4 | 49.4 | 49.4 | 50.2 |
| Defocus Range (µm) | -0.8 to -2.0 | -0.8 to -2.0 | - 0.5 to - 1.8 | -0.8 to -1.6 | -0.8 to -1.6 | -0.8 to -1.6 | -0.8 to -1.6 | -1.0 to -1.8 |
| Pixel Size (Å) | 0.8015 | 0.8015 | 1.045 | 1.045 | 1.045 | 1.045 | 1.045 | 1.150 |
| Imposed Symmetry | C3 | C3 | C3 | C3 | C3 | C3 | C3 | C3 |
| Number of Micrographs | 3798 | 3798 | 4998 | 5946 | 5297 | 5297 | 5297 | 4044 |
| Final Particle Number (Symmetry Expanded) | 23,695 | 51,733 | 23,375 | 24,189 | 12,893 | 140,230 | 80,633 | 33,100 |
| Map Resolution (Å) | 3.8 | 3.3 | 3.3 | 4.4 | 4.6 | 3.8 | 4.0 | 5.8 |
| FSC Threshold | 0.143 | 0.143 | 0.143 | 0.143 | 0.143 | 0.143 | 0.143 | 0.143 |
| Map Sharpening B-factor ($Å^2$) | -79.3 | -83.6 | -66.5 | -118.9 | -128.6 | -117.1 | -124.0 | -147.9 |
| **Model Refinement and Validation** | | | | | | | | |
| Total Residues | 2247 | 2243 | 3118 | 2245 | 2527 | 2099 | 2109 | - |
| Amino Acids | 2170 | 2164 | 3044 | 2156 | 2428 | 1995 | 2006 | - |
| Carbohydrates | 77 | 84 | 74 | 89 | 99 | 104 | 103 | - |
| RMSD Bonds | 0.004 | 0.006 | 0.005 | 0.005 | 0.006 | 0.005 | 0.004 | - |
| RMSD Angles | 0.71 | 0.97 | 0.85 | 0.86 | 0.845 | 0.71 | 0.67 | - |
| **Ramachandran** | | | | | | | | |
| Outliers (%) | 0 | 0 | 0 | 0 | 0 | 0 | 0 | - |
| Allowed (%) | 2.9 | 1.1 | 3.4 | 3.2 | 4.3 | 2.6 | 2.8 | - |
| Favored (%) | 97.1 | 98.9 | 96.6 | 96.8 | 95.7 | 97.4 | 97.2 | - |
| Rotamer Outliers (%) | 0 | 0 | 0.13 | 0.06 | 0 | 0 | 0 | - |
| Cβ Outliers (%) | 0 | 0 | 0 | 0 | 0 | 0 | 0 | - |
| Clash Score | 3.50 | 2.04 | 4.38 | 4.16 | 4.80 | 4.48 | 2.21 | - |
| Molprobity Score | 1.30 | 0.97 | 1.44 | 1.40 | 1.54 | 1.34 | 1.14 | - |
| Map Correlation Coefficient | 0.83 | 0.86 | 0.83 | 0.77 | 0.74 | 0.80 | 0.80 | - |
| FSC Model (0/0.143/0.5) | 3.7/3.8/3.9 | 3.2/3.3/3.4 | 3.2/3.3/3.4 | 4.2/4.3/4.5 | 4.4/4.5/4.7 | 3.6/3.8/4.0 | 3.9/3.9/4.1 | - |
| EMRinger Score | 3.18 | 4.18 | 3.44 | 1.78 | 0.79 | 2.73 | 2.15 | - |

### Table S2.

**Supplemental Table 2. Electron Microscopy Data Bank deposition information for nsEMPEM models.** Maps are accessible at emdataresource.org using the listed codes. Additional maps may be found on the "Download" tab for each entry.

| EMDB Code | SOSIP | Animal ID | Timepoint | Map | Polyclonal Antibody Epitopes |
|---|---|---|---|---|---|
| **EMD-43895** | 16055.v8.3 | r2464 | W6 | Main Map | Base |
| **EMD-43896** | 16055.v8.3 | r2464 | W22 | Main Map | Base, N611, V2 |
| | | | | Main Map | N611/N625, V1/V2/V3 |
| **EMD-43909** | 16055.v8.3 | r2463 | W22 | Additional Map 1 | Base, V1/V2/V3 |
| | | | | Additional Map 2 | C3/V5, V1/V2/V3 |
| **EMD-43910** | CH505/BG505 SOSIP.v8.1 | r2474 | W6 | Main Map | Base |
| | | | | Main Map | Base, C3/V5, V1/V3, V2, Anti-IC |
| **EMD-43911** | CH505/BG505 SOSIP.v8.1 | r2474 | W26 | Additional Map 1 | gp120-Interface, Base, C3/V5, V2 |
| | | | | Additional Map 2 | Base, N611, V2, C3/V5 |
| **EMD-43915** | B41 SOSIP.v4.1 | r1646 | W4 | Main Map | N611 |
| | | | | Additional Map 1 | Base |
| **EMD-43916** | B41 SOSIP.v4.1 | r1646 | W6 | Main Map | Base, N611 |
| **EMD-43917** | B41 SOSIP.v4.1 | r1646 | W22 | Main Map | N241, Base, Anti-IC |
| | | | | Additional Map 1 | N241, Base, N611 |
| **EMD-43918** | BG505 SOSIP.v5.2 (N241/N289) | Rh.33203 | W10 | Main Map | N289, Base |
| **EMD-43919** | BG505 SOSIP.v5.2 (N241/N289) | Rh.33203 | W26 | Main Map | gp120-Interface, N289, Base, C3/V5 |
| **EMD-44076** | BG505 SOSIP.v5.2 (N241/N289) | Rh.33203 | W38 | Main Map | gp120-Interface, Base, C3/V5, Anti-IC |
| | | | | Additional Map 1 | gp120-Interface, N289, Base, C3/V5 |
